# Three-dimensional mobility and muscle attachments in the pectoral limb of the Triassic cynodont *Massetognathus pascuali*

**DOI:** 10.1101/166587

**Authors:** Phil H. Lai, Andrew A. Biewener, Stephanie E. Pierce

## Abstract

The musculoskeletal configuration of the mammalian pectoral limb has been heralded as a key anatomical feature leading to the adaptive radiation of mammals, but limb function in the cynodont outgroup remains unresolved. Conflicting reconstructions of abducted and adducted posture are based on mutually-incompatible interpretations of ambiguous osteology. We reconstruct the pectoral limb of the Triassic non-mammalian cynodont *Massetognathus pascuali* in three dimensions, by combining skeletal morphology from micro-computed tomography with muscle anatomy from an extended extant phylogenetic bracket. Conservative tests of maximum range of motion suggest a degree of girdle mobility, as well as substantial freedom at the shoulder and the elbow joints. The glenoid fossa supports a neutral pose in which the distal end of the humerus points 45° posterolaterally from the body wall, intermediate between classically “sprawling” and “parasagittal” limb postures. *Massetognathus* is reconstructed as having a near-mammalian complement of shoulder muscles, including an incipient rotator cuff (m. subscapularis, m. infraspinatus, m. supraspinatus, and m. teres minor). Based on close inspection of the morphology of the glenoid fossa, we hypothesize a posture-driven scenario for the evolution of the therian ball-and-socket shoulder joint. The musculoskeletal reconstruction presented here provides the anatomical scaffolding for more detailed examination of locomotor evolution in the precursors to mammals.

## INTRODUCTION

Today’s mammals inhabit disparate ecological niches, comprising cursorial, fossorial, aquatic, and even volant forms (Vaughan et al. 2013; Hildebrand, 1989; Fischer et al. 2002). These varied lifestyles are supported by modifications of the pectoral limb into anatomical structures as diverse as wings and flippers. The evolution of the mammalian-style pectoral limb—mobile scapula, ball-and-socket glenohumeral joint, “parasagittal” limb posture—has been suggested as a key innovation leading to the adaptive radiation of the clade (Polly, 2007). Morphological diversification of this anatomical module began early on in mammalian evolution (Meng et al. 2006; Ji et al. 2006), predating the emergence of the crown group (Luo, 2007). Accordingly, interpreting morphological and functional transformation of the pectoral limb in the sister group to mammals is key to understanding their remarkable success.

The non-mammalian cynodonts offer a glimpse at an intermediate stage in mammalian locomotor evolution. The osteology of the cynodont pectoral girdle and forelimb is well known from the fossil record, and does not appear to have been particularly disparate; in a series of papers, Jenkins (1970a, 1971a) synthesized a number of descriptions and posited that most cynodonts shared a common appendicular morphology, and presumably similar locomotor behaviors. In contrast to the hip articulation, where a socket-like acetabulum clearly circumscribed range of motion (Jenkins, 1971a), the cynodont gleno-humeral joint possessed the relatively unconstrained, hemisellar architecture on which late Permian archosaurs, lepidosaurs, and synapsids converged (Jenkins, 1993)—the typical mammalian ball-and-socket articulation did not appear until the Jurassic theriimorphs (Ji et al. 1999; Luo, 2015). Multiple reconstructions of the cynodont pectoral limb have been advanced, drawing on skeletal morphology (Watson, 1917; Jenkins, 1970b, 1971a; Kemp, 1980a, 1980b; Oliveira & Schultz, 2016) as well as muscle anatomy as inferred from osteology and homology to extant taxa (Gregory & Camp, 1918; Romer, 1922).

Two competing hypotheses of cynodont posture and locomotion have emerged, with discrepancies centered on divergent interpretations of shoulder mobility, and the position occupied along the classic sprawling-to-upright continuum of tetrapod posture (Reilly & Elias, 1998; Gatesy, 1991). A cornerstone of the upright, or adducted postural view, is Jenkins’ work on *Massetognathus pascuali*, a traversodontid cynodont from the Triassic Chañares Formation of Argentina (Romer, 1967; Jenkins, 1970b). As a member of Cynognathia, the sister group to the probainognathians that gave rise to mammals (Ruta et al. 2013), *Massetognathus* represents one of the last steps on the mammal stem, as well as a reasonable exemplar of early Mesozoic cynodont anatomy (Liu & Olsen, 2010). Based on the postcranial skeleton of *Massetognathus*, Jenkins advanced a two-dimensional reconstruction in a crouched, adducted pose with posteriorly-directed elbows, reminiscent of a small, short-limbed therian (Jenkins, 1970b). Working from cynognathian (*Cynognathus*) and probainognathian (*Trucidocynodon*) material, Watson (1917) and Oliveira and Schultz (2016) arrived at similarly therian-like interpretations of posture and locomotion across eucynodonts.

On the other hand, Kemp’s ( 1980a, 1980b) reconstructions of the basal Late Permian cynodont *Procynosuchus* and the Middle Triassic traversodontid *Luangwa* depicted the sprawling, abducted posture thought to be plesiomorphic for amniotes. The humerus is held perpendicular to the animal’s sagittal plane, and the main stride component is furnished by protraction and retraction of the humerus around a dorsoventral axis. Kemp posited that the basic structure and function of the forelimb remained unchanged between the Permian and Triassic cynodonts, and that the limb and girdle transformations leading to adducted posture were restricted to later, more crownward taxa. The close phylogenetic relationship between *Massetognathus* and *Luangwa* (Liu & Abdala, 2014) means that we currently have a reconstruction with abducted posture in one traversodontid, and adducted posture in another. The equivocal osteology of the cynodont pectoral girdle has so far precluded consensus on forelimb function, hindering a deeper understanding of locomotor evolution in this important clade.

Here we revisit cynodont forelimb morphology and function using modern computational methods to add a third dimension (3D) to this classic problem. Using digital models of fossil material derived from micro-computed tomography (µCT), we interactively assess articular function at the shoulder and elbow joints (e.g. Pierce et al. 2012; Nyakatura et al. 2015). Further, we reconstruct the origins and insertions of the shoulder musculature using an updated, extended extant phylogenetic bracket, and map them onto the 3D pectoral limb skeleton of *Massetognathus*. The result is a robust, three-dimensional reconstruction that will form the basis of future biomechanical analyses using musculoskeletal modeling techniques (e.g. Bates & Schachner, 2012; Hutchinson et al. 2005; Hutchinson et al. 2015) to probe the link between skeletal motion and muscle function.

## MATERIALS AND METHODS

### µCT scanning and segmentation

A nodule containing the nearly-complete, articulated remains of *Massetognathus pascuali* (MCZ 3691) was scanned using a Nikon Metrology (X-Tek) HMXST225 MicroCT unit located at Harvard University’s Center for Nanoscale Systems. Scanning parameters were 175kV 46µA, with a 0.01mm copper filter and a final voxel size of 127.22 µm. The µCT data were imported into Mimics v18 (Materialise NV, Leuven, Belgium) for segmentation. Pectoral girdle (interclavicle, clavicles, scapulocoracoids) and forelimb (humeri, radii, ulnae) skeletal elements were identified and assigned individual masks, from which high-resolution 3D meshes were computed and exported for smoothing and repair (Fig. 1).

**Figure 1.**
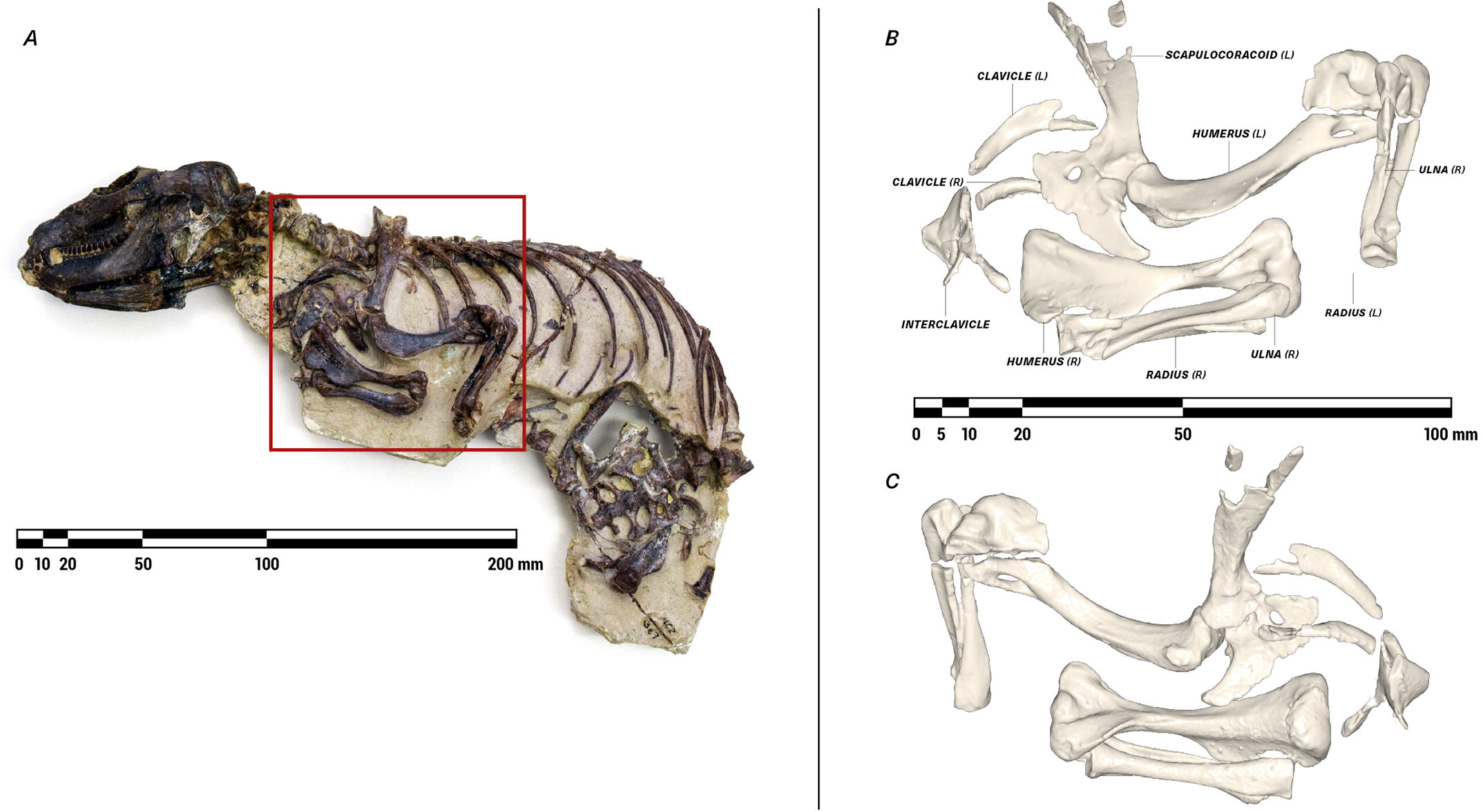
Nodule containing the articulated remains of *Massetognathus pascuali* (MCZVP 3691) (A), with lateral (B) and medial (C) views of pectoral limb 3D surface models, prior to mesh refinement and repair. MCZVP, Museum of Comparative Zoology, Department of Vertebrate Paleontology, Harvard.

### Bone repair

The pectoral limb exhibited no overall distortion, but certain bones had suffered taphonomic fragmentation, necessitating repairs to their digital models. Fragments were manually aligned, using contralateral elements as reference. In the case of the humerus, radius, and clavicle, damage to opposing ends of the left- and right-side elements was remedied by taking the intact end of one element, mirroring it, and grafting it onto the opposing end of its counterpart. Small gaps were filled using the “Wrap” and “Surface Reconstruction” algorithms in 3-Matic v10 (Materialise NV, Leuven, Belgium), while larger breaks were bridged and smoothed over using digital sculpting tools in Autodesk Mudbox (Autodesk, Inc., San Rafael, CA, USA). As we were unable to locate all of the fragments of the interclavicle, we modeled missing segments based on the preserved cranial portion, other interclavicles in the MCZ collections, and Jenkins’ (1970b, 1971a) description of the same element in other cynodonts.

The repaired bone meshes were smoothed and re-wrapped in MeshLab (ISTI-CNR, Pisa, Italy), to eliminate artifacts introduced in scanning and segmentation, while preserving potentially informative surface texture. To reduce noise, we performed a Poisson surface reconstruction (Hoppe, 2008), which takes the vertex coordinates of the original mesh and outputs an optimized, re-triangulated mesh. We then applied a single Laplacian smoothing step (Field, 1988) to correct any remaining polygonal irregularities, and exported the finished meshes in Wavefront .OBJ format (Wavefront Technologies, Santa Barbara, CA, USA) for examination and assembly.

### Re-articulation and rigging

Using 3DS Max (Autodesk, Inc., San Rafael, CA, USA), centers of rotation for the acromio-clavicular, humero-radial, and humero-ulnar joints were determined by fitting spherical primitives to their opposing articular surfaces and then superimposing the spheres’ centroids in 3D space ("2). A cylinder was used to model the planar clavo-interclavicular joint, and an ellipsoid of aspect ratio 24:13 was used to model the gleno-humeral joint. Thus articulated, the pectoral girdle and forelimb skeleton were organized as a kinematic hierarchy, wherein each bone was subordinated to the reference frame of its proximal neighbor, and inherited all rotations and translations applied to the latter.

A local joint coordinate system (JCS) (Grood & Suntay, 1983) was defined for each articulation, with joint axes positioned to reflect anatomically-informative rotations. Axes were oriented following the XYZ rotation order convention, with Z capturing the axis of greatest expected mobility and X the least (Brainerd et al. 2010).

### Range of motion testing

Limb joint range of motion has been shown to be sensitive to assumptions of intra-joint spacing (Arnold et al. 2014; Nyakatura et al. 2015; Pierce et al. 2012). To circumscribe this, we established an articular cartilage thickness of 0.25 mm based on relationships for mammalian articular cartilage (Simon, 1970). Assuming equal cartilage thickness on opposing articular surfaces, we modeled every joint with a total joint space of 0.50 mm. This value is supported by the *in situ* spacing between limb elements in the fossil specimen, and yielded good agreement between the curvature of opposing articular surfaces.

Osteological limits to joint motion were assessed by rotating the distal element of each joint until it collided with another bony surface, and repeating in the opposite direction to give total osteological range of motion around each axis. The clavo-interclavicular, acromio-clavicular, humero-radial, and humero-ulnar joints were tested only in rotation, but the unusual morphology of the gleno-humeral joint has been suggested to support coordinated translation and rotation, in the form of sliding (Jenkins, 1971a) or rolling (Kemp, 1980b) kinematics. Accordingly, we opted to compare purely rotational range of motion at this joint against combined translation-rotation. To do so, we imported the *Massetognathus* model into SIMM (Software for Interactive Musculoskeletal Modeling: Delp & Loan, 1995) and defined kinematic functions linking rotations around X, Y and Z (Fig. 2) with translations along those same axes. The functions were tuned to maintain a constant 0.5mm offset between the humeral head and the glenoid fossa, as determined via collision between a second, larger ellipsoid primitive and the surface of the glenoid fossa. Due to the unconstrained morphology of the glenoid fossa, we found that extremes of gleno-humeral rotation could result in disarticulation of the joint well before a collisional limit was reached. To establish reasonable physiological limits for this joint, we defined a secondary constraint criterion of 50% humeral head contact with the glenoid fossa. Results of range of motion testing are given in Table 1.

**Figure 2.**
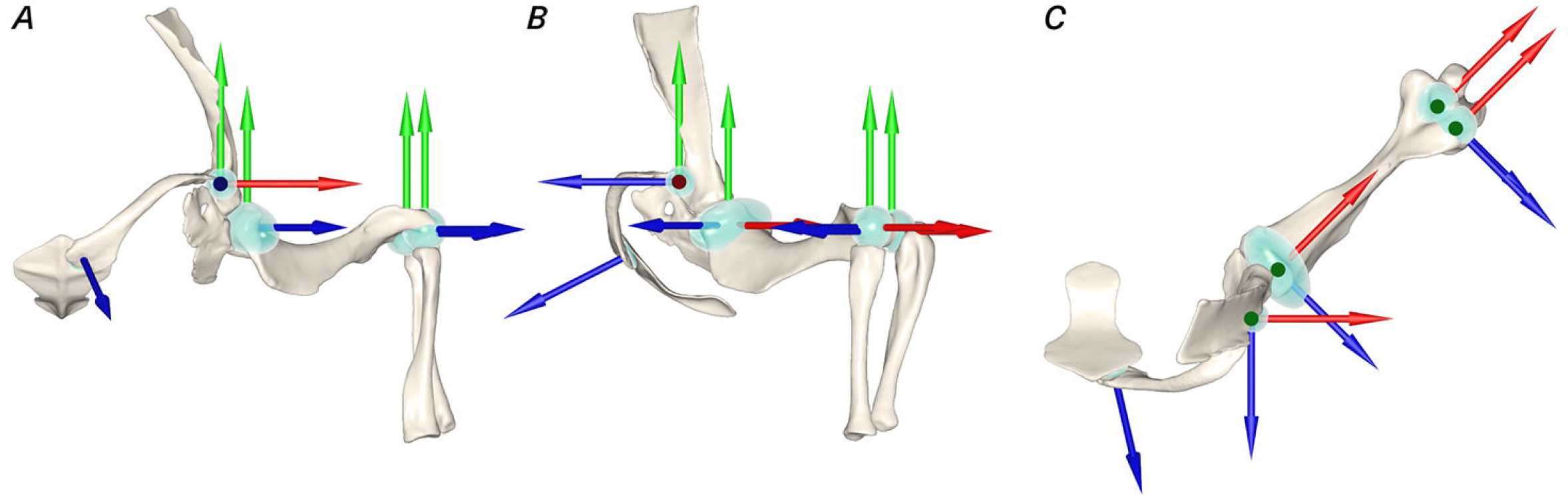
Cranial (A), lateral (B), and dorsal (C) views of the articulated left-side pectoral limb of *M. pascuali*, showing rotational axes and primitives used to determine centers of rotation. X, Y, and Z axes as labeled in Table 1. Axis colors: X-Red/Y-Green/Z-Blue.

**Table 1.**
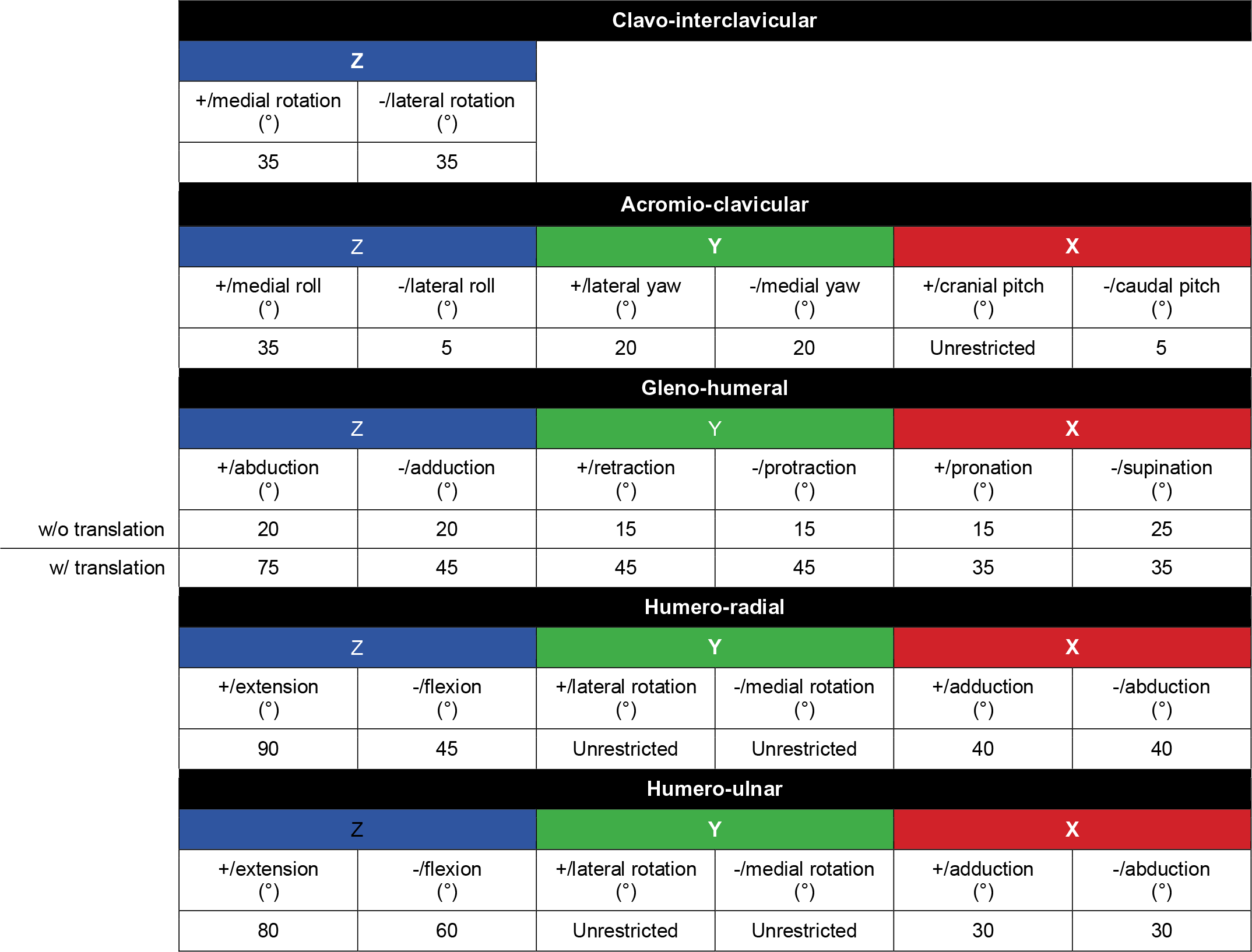
Pectoral limb joint range of motion in the cynodont *Massetognathus pascuali*. All measurements are made using 0.5mm joint space (see text for details).

A neutral reference pose was defined, allowing joint rotations to be repeatedly measured and compared (Brainerd et al. 2010; Gatesy et al. 2010). The gleno-humeral joint was rotated to the center of its measured ranges of motion (Fig. 3). Doing so placed the approximate centroids of the glenoid fossa and the humeral head in apposition, as has been hypothesized to best approximate *in vivo* utilization of joint surfaces during locomotion (Fischer, 1994). The humero-radial and humero-ulnar joints were then rotated to orient the antebrachium normal to the “substrate”.

**Figure 3.**
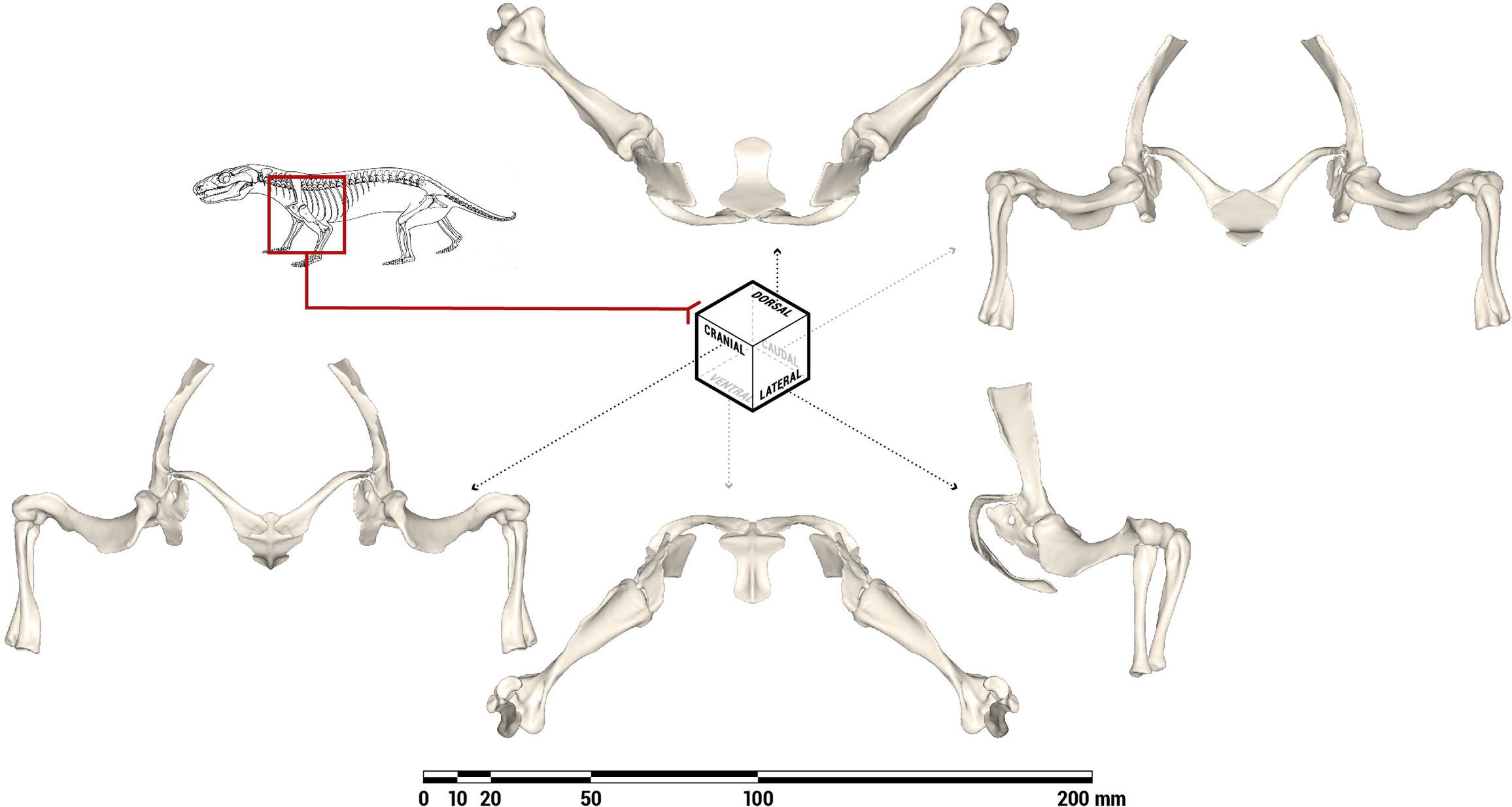
Orthographic views of the pectoral limb of *M. pascuali* in an anatomically-neutral reference pose (not in vivo posture), with all joints rotated to the centers of their measured ranges of motion. The bones depicted comprise the bilateral scapulocoracoids, humeri, ulnae, and radii, as well as the median interclavicle. Line drawing of *Massetognathus* adapted from Figure 9 in Jenkins (1970b).

### Shoulder muscle reconstruction

Prior reconstructions of cynodont musculature have used monotremes (Gregory & Camp, 1918) as well as saurians and therians (Romer, 1922; Jenkins, 1971b) as bookends to examine conservation and transformation in muscle anatomy through mammal evolution. Following this practice, we defined an extended Extant Phylogenetic Bracket (Witmer, 1995) encompassing a range of amniotes, and using salamanders as an outgroup (see Table 2 for the full list of taxa). Due to the rarity of articulated cynodont carpi and the paucity of well-delineated attachment sites on the radius and ulna, past studies have largely focused on muscles crossing the shoulder joint. In the interest of parsimony and repeatability, we follow suit in limiting our set of muscles under consideration to only those that span the gleno-humeral joint. This metric excludes the extrinsic scapular muscles (such as m. serratus anterior, m. trapezius, and mm. rhomboidei) that originate on the axial skeleton and insert on the shoulder girdle, as well as the various flexors, extensors, pronators, and supinators that originate on the distal humerus and actuate the distal forelimb. M. latissimus dorsi and mm. pectorales have insertions on the humerus, and were included in the analysis despite their origins on the axial skeleton, due to their presumed first-order action on the brachium.

**Table 2.**
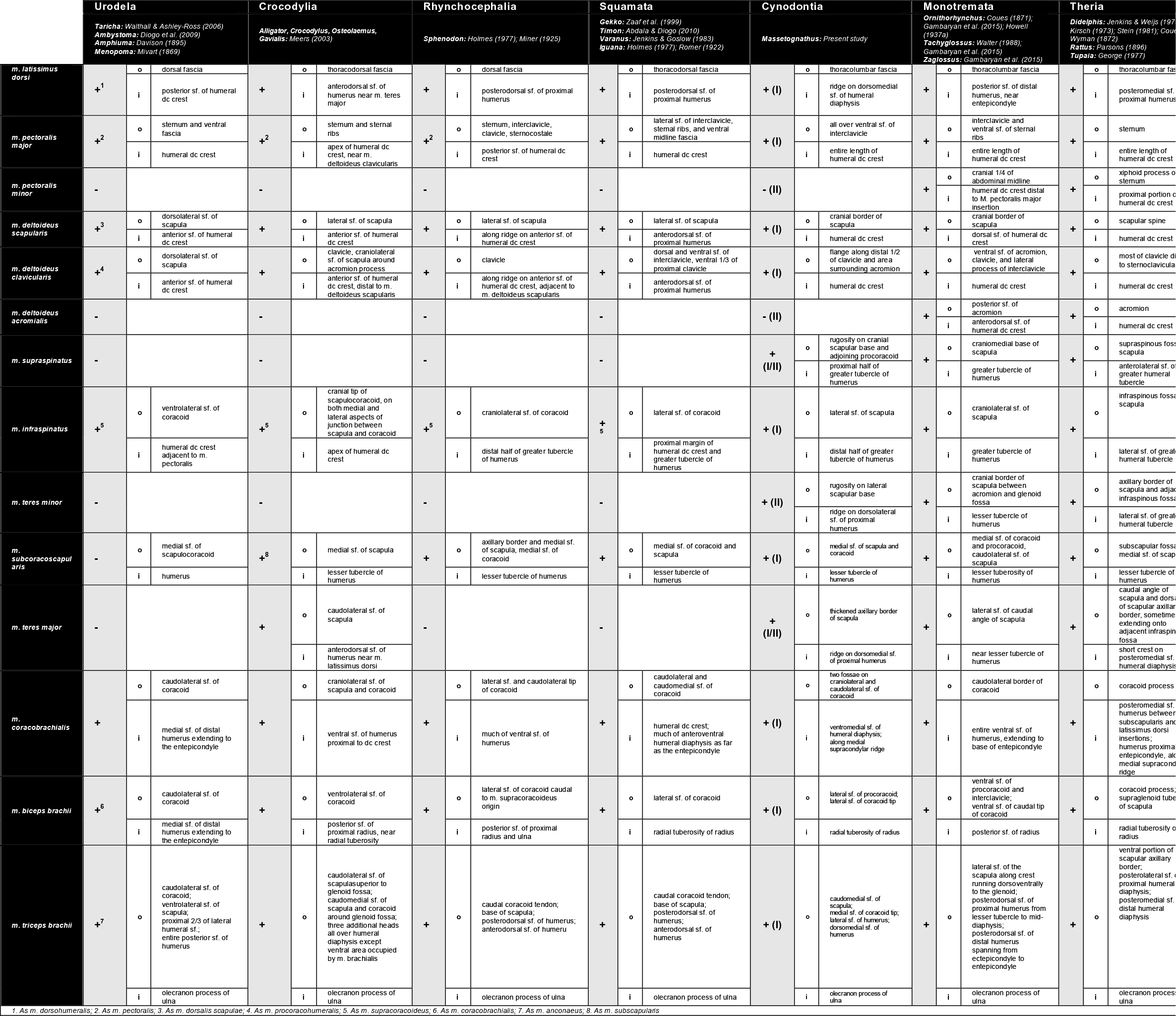
Extant Phylogenetic Bracket with muscle attachments and homologies. Abbreviations: sf., surface; dc, deltopectoral crest

Muscle origins and insertions were taken from the primary literature (see Table 2 for list of references), and reconstructed as likely present in *Massetognathus* (level I inference *sensu* Witmer, 1995) or possibly present (level II inference). None of the muscles under consideration were determined to be likely absent (level III inference). Bryant and Seymour’s (1990) study of carnivorans was used as a reference for muscle attachment type (direct/fleshy, aponeurotic, or tendinous). Absence of clear osteological correlates was not considered grounds for elimination, as muscles inserting directly into periosteum may not leave a mineralized scar (Bryant & Seymour, 1990). In the absence of bony scars, we placed attachments on homologous regions of the bone (Holliday, 2009). Muscles were homologized with reference to Abdala and Diogo (2010) and Diogo et al. (2009). Reconstructed muscle attachment areas were then digitally painted onto the 3D bone meshes using Autodesk Mudbox, for visualization and comparison.

## RESULTS

### Neutral reference pose

Digitally reassembling the pectoral limb into a neutral reference pose (Fig. 3) places the diaphysis of the humerus at approximately 45° to the animal’s body wall, with the proximal and distal articular surfaces in roughly the same horizontal plane. The anteriorly-oriented scapulocoracoid, posterolateral placement of the glenoid fossa, and caudally-pointing elbows of this pose are broadly consistent with Watson’s (1917), Jenkins’ (1971a), and Kemp’s (1980a) reconstructions of traversodontid pectoral girdles/limbs.

Jenkins (1971a) contested Watson’s (1917) reconstruction of the cynodont pectoral girdle, arguing that angling the scapulocoracoids medially and cranially would force the humerus into a mechanically untenable posture while compromising the weight-bearing suspensory function of the extrinsic scapular musculature. Jenkins’ reconstruction orients the scapulocoracoids more vertically and tilts them outward from the midline; as a result, the glenoid fossae are directed more ventrally than in Watson’s reconstruction. Based on our 3D reconstruction, the geometry of the clavicles and interclavicle constrain the scapulocoracoids such that the glenoid fossae must be oriented posterolaterally and slightly ventrally, as in Jenkins’ reconstruction.

### Joint range of motion

As reconstructed here, the pectoral limb of *Massetognathus pascuali* has one possible degree of rotational freedom (DOF) at the clavo-interclavicular joint, up to three at the acromio-clavicular joint, and three each at the gleno-humeral, humero-radial, and humero-ulnar joints, for a total of 13 DOF. Table 1 presents measured ranges of motion at each joint.

The clavo-interclavicular articulation lacks the extensive, rigid transverse overlap seen in monotremes (Luo, 2015), and there are no known instances of synostosis between these elements in *Massetognathus*. The planar geometry of the articular facets limits assessment of mobility at this joint to medial and lateral rotation of the clavicle around an axis normal to the interclavicle (Figure 2) (Kemp, 1980b). Clavicular elevation-depression and long-axis rotation are improbable, as such movements would require compressive deformation of a substantial thickness of soft tissue around the joint.

Some degree of mobility at the acromio-clavicular articulation has been hypothesized (Kemp, 1980b), but the extent of this is difficult to assess in our 3D model of *Massetognathus* as the acromion is small and possibly incomplete. Although the minimally-projecting acromion observed in MCZ 3691 is consistent with Jenkins’ (1971a) and Kemp’s (1980a) descriptions of eucynodont scapulocoracoids, Liu (2007) documented several traversodontid scapulocoracoids with more prominent acromions, including a juvenile *Massetognathus pascuali*. The distal tip of the clavicle likely presents a shallowly, concave articular surface that is somewhat congruent with the medial surface of the presumptive acromion. We cautiously propose that some sliding or translational motion may have been possible between the clavicle and the acromion, allowing pitch, roll, and yaw rotations around the acromio-clavicular joint, subject to soft-tissue constraints.

The glenoid fossa is dorsoventrally concave and anteroposteriorly convex, resembling one half of a sellar joint. The humeral head has an approximately ellipsoidal morphology, with the major axis running between the greater and lesser tubercles. Rotation appears to be somewhat restricted in the absence of translation, totalling 40° in abduction-adduction, 30° in retraction-protraction, and 40° in pronation-supination. By contrast, we found greatly increased mobility around all three rotational axes when allowing for translation, suggesting—in line with Jenkins (1971a) and Kemp (1980b)—that the gleno-humeral articulation is a six degree of freedom joint.

Moving distally to the elbow, we measured comparable amounts of total flexion-extension for the humero-radial and humero-ulnar joints (135° for the former, 140° for the latter). Both the radius and the ulna are capable of some amount of abduction-adduction (80° for the radius, 60° for the ulna), and long-axis rotation of each of these bones is unrestricted with reference to the neutral pose. In life, interosseous ligaments and pronator muscles running between the radius and ulna would likely have constrained independent movement of these two bones, while permitting coordinated pronation and supination within the maximum osteological ranges established here.

### Muscle reconstruction

The full set of muscles and taxa considered is given in Table 2, along with primary literature references. A total of 12 muscles were reconstructed for *Massetognathus pascuali*. Individual muscles and their attachments are discussed in the text below.

### M. latissimus dorsi (Fig. 4)

M. latissimus dorsi appears to be plesiomorphic for tetrapods (Romer, 1924), and is present across the phylogenetic bracket (Table 2). This muscle originates aponeurotically from the dorsal and thoracodorsal fascia, and sometimes takes multiple costal origins as well (Diogo et al., 2009). It inserts on the proximodorsal surface of the humeral deltopectoral crest in all cases except for monotremes, where the insertion follows the deltopectoral crest distally to terminate on the entepicondyle (Gambaryan et al. 2015). M. latissimus dorsi in non-mammalian cynodonts is a level I inference, given its presence on both sides of the bracket. This muscle inserts adjacent to m. teres major on a linear area running parallel to the long axis of the humerus (Fig. 4). Based on *Cynognathus*, Jenkins (1971a) situated the insertion of the cynodont m. latissimus dorsi on a ridge running obliquely across the dorsal surface of the humerus. He did not identify a corresponding ridge in *Thrinaxodon* or *Massetognathus*, and we are unable to locate this feature on close examination of the latter. Instead, we follow Watson (1917), Romer (1922) and Kemp (1980a) in reconstructing a linear insertion for m. latissimus dorsi running proximodistally along the dorsomedial surface of the humerus, terminating on a tuberosity just proximal to the midpoint of the diaphysis.

**Figure 4.**
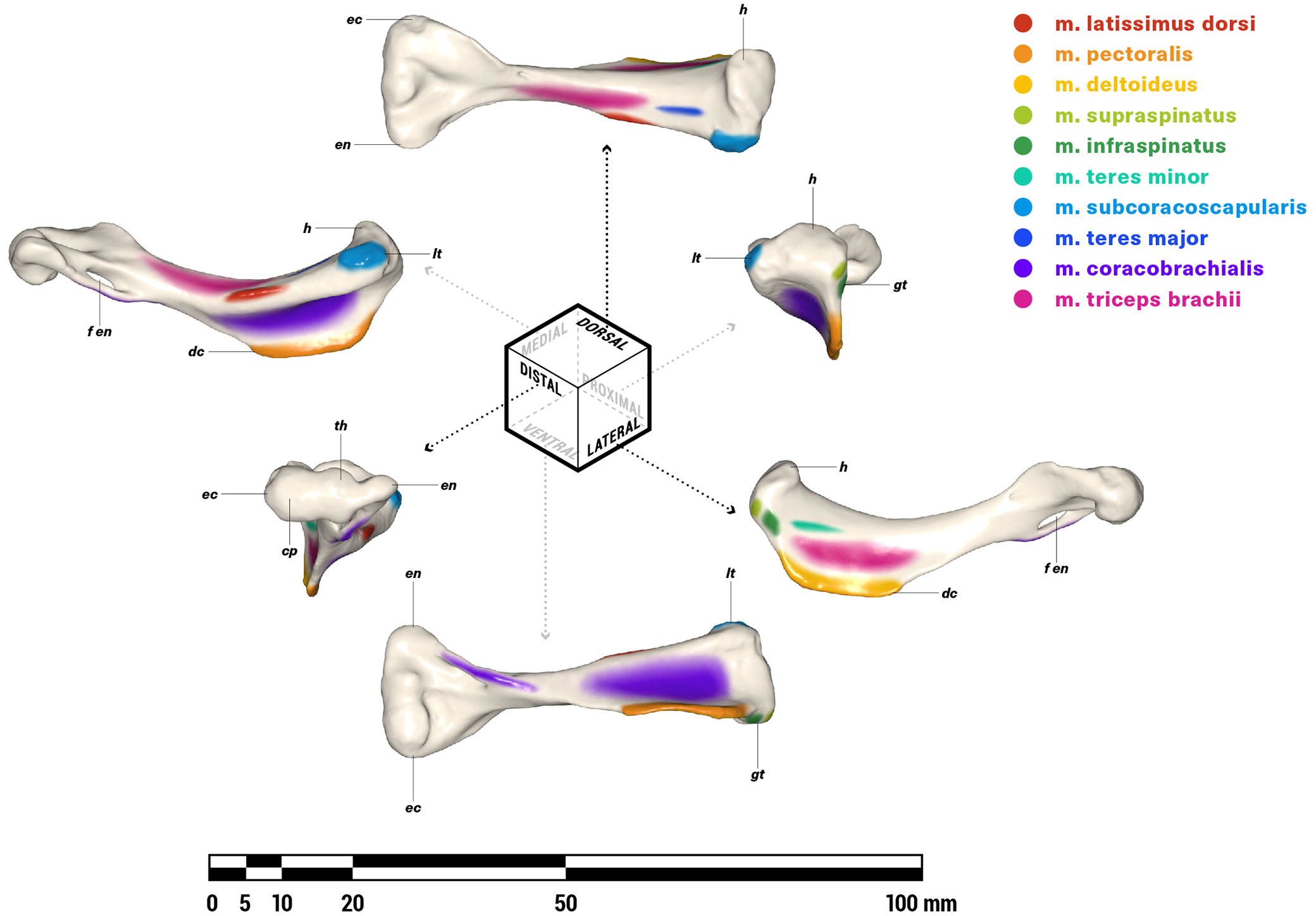
Orthographic views of the left humerus of *M. pascuali*, with reconstructed muscle origins/insertions. Abbreviations: cp, capitulum; dc, deltopectoral crest; ec, ectepicondyle; en, entepicondyle; f en, entepicondylar foramen; gt, greater tubercle; h, humeral head; lt, lesser tubercle; th, trochlea. Reconstructed muscles are listed in legend.

### M. pectoralis (Figs. 4, 5)

M. pectoralis is present in all extant tetrapods (Table 2). While non-mammals may have multiple m. pectoralis heads (Jenkins & Goslow, 1983), the mammalian pectoralis complex comprises a cranial, superficial pectoralis major and a caudal, deeper pectoralis minor (Jenkins & Weijs, 1979). In most mammals, these muscles insert together along the length of the humeral deltopectoral crest, but in some taxa (including humans) the pectoralis minor inserts separately on the coracoid process of the scapula. This is certainly a derived condition, and for the purposes of this study we will consider m. pectoralis as a single functional unit, without major or minor divisions. Originating from the ventral midline of all tetrapods on (where present) the interclavicle, sternal series, and sometimes the medial ends of the costal cartilages, m. pectoralis inserts on the posteroventral surface or apex of the deltopectoral crest in all cases. *Massetognathus* possesses a prominent deltopectoral crest running slightly more than halfway along the humeral diaphysis. A “cruciate” interclavicle is plesiomorphic for synapsids (Jenkins, 1971a), and was considered by Romer (1940) to give origin to a pectoralis complex via the paired fossae on the posterior ramus. Relative to its length, the interclavicle of *Massetognathus* is considerably broader mediolaterally than that of pelycosaurs, with a well-marked posterior ramus and ridge that may represent an expanded attachment for m. pectoralis (Romer, 1940; Jenkins, 1971a). We reconstruct m. pectoralis as a level I inference, originating all over the lateral surfaces of the posterior process, across the fossae on the posterior ramus, and possibly also on the caudally-facing surfaces of the lateral ridge (Fig. 5). M. pectoralis inserts as an aponeurosis on the posteroventral surface of the deltopectoral crest of the humerus, spanning its proximodistal length (Fig. 4).

**Figure 5.**
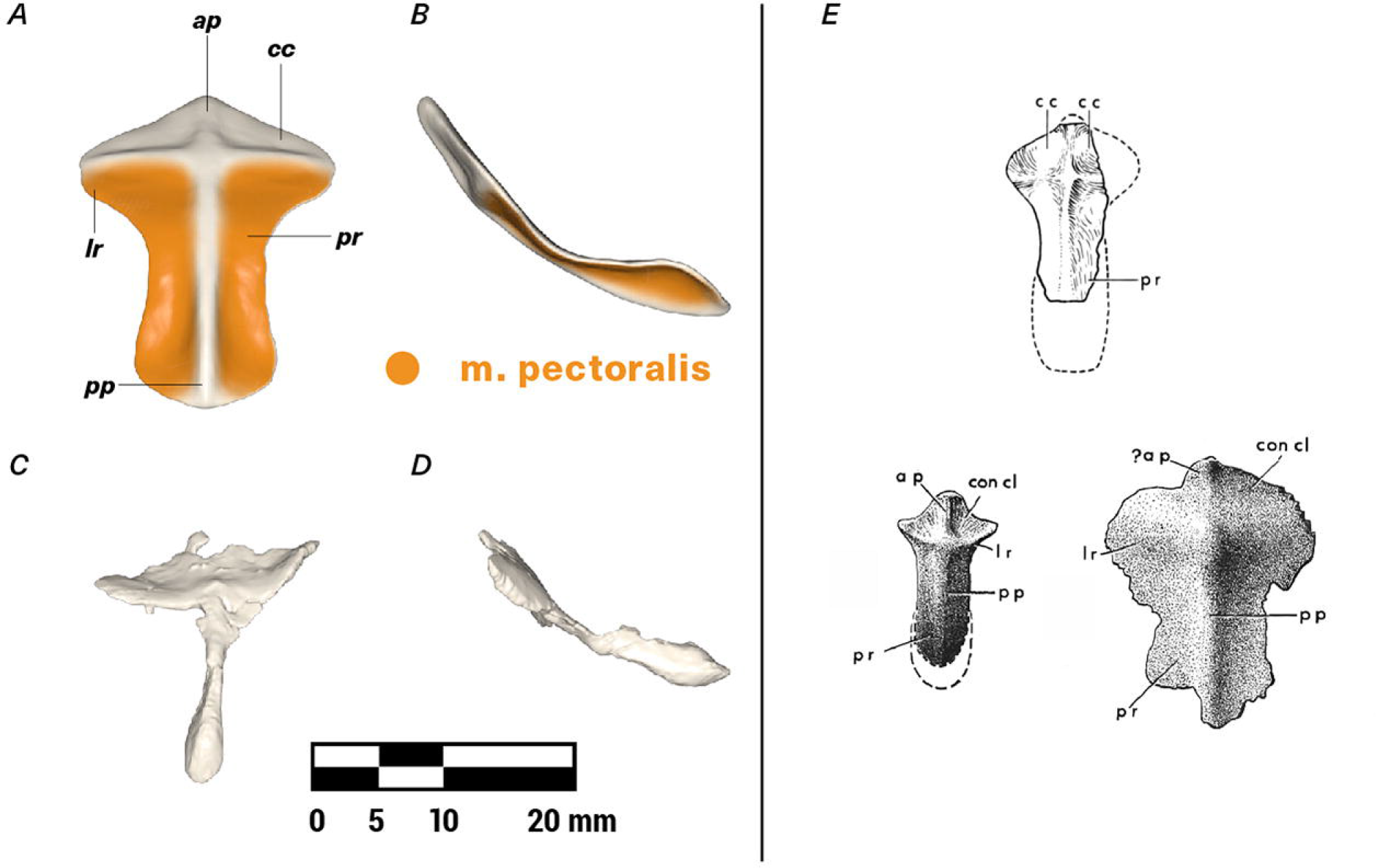
Repaired (A, B) and original (C, D) interclavicle of *M. pascuali* in ventral (A, C) and lateral (B, D) views, with reconstructed muscle origins/insertions. Reference images (E) adapted from Jenkins (1970b) (top, *Massetognathus*) and Jenkins (1971a) (bottom left *Thrinaxodon*, bottom right unidentified cynodont.) Abbreviations follow Jenkins (1971a): ap, anterior ridge; cc/con cl, concavity for clavicle articulation; lr, lateral ridge; pp, posterior ridge; pr, posterior ramus. Reconstructed muscles are listed in legend.

### M. deltoideus scapularis and m. deltoideus clavicularis (Figs. 4, 6, 7)

M. deltoideus is present in all tetrapods as a scapular division (m. deltoideus scapularis) and a clavicular division (m. deltoideus clavicularis), with mammals gaining an additional acromial division (m. deltoideus acromialis) (Table 2, homology follows Diogo et al. 2009). The acromion appears to be variably-developed in *Massetognathus* and other cynodonts, and offers no obvious site for muscle attachment (Jenkins, 1971a; Kemp, 1978); accordingly, we have reconstructed *Massetognathus* with only the scapular and clavicular heads common to all tetrapods.

Like monotremes, the cynodont scapula has a strongly reflected cranial border (rcb, Fig. 7), which is probably homologous to the therian scapular spine (Romer, 1922; Jenkins, 1971a; Kemp, 1980a, Gambaryan et al. 2015). The caudally-facing surface of this border is the likely site of origin for m. deltoideus scapularis, in agreement with Gregory and Camp (1918), Romer (1922), and Jenkins (1971a), but *contra* Kemp (1980a, 1980b), who attributed a broader origin to m. deltoideus, covering much of the lateral surface of the scapula in addition to the reflected border. In living lepidosaurs and crocodylians, m. deltoideus scapularis takes origin from the cranial or craniodorsal portions of the lateral scapular surface. The presence of a pronounced reflected cranial border would functionally divide an m. deltoideus scapularis spanning the entire lateral surface of the scapula into a posteriorly-facing portion and a laterally-facing portion, with uncertain consequences for its resultant line of action. In the absence of any instances across the phylogenetic bracket of such a functional division, we consider an m. deltoideus scapularis origin restricted to the reflected cranial border more biomechanically plausible. As in all other tetrapods (Table 2), the cynodont m. deltoideus scapularis inserts in conjunction with m. deltoideus clavicularis, on the anterodorsal surface of the humeral deltopectoral crest (Fig. 4).

**Figure 6.**
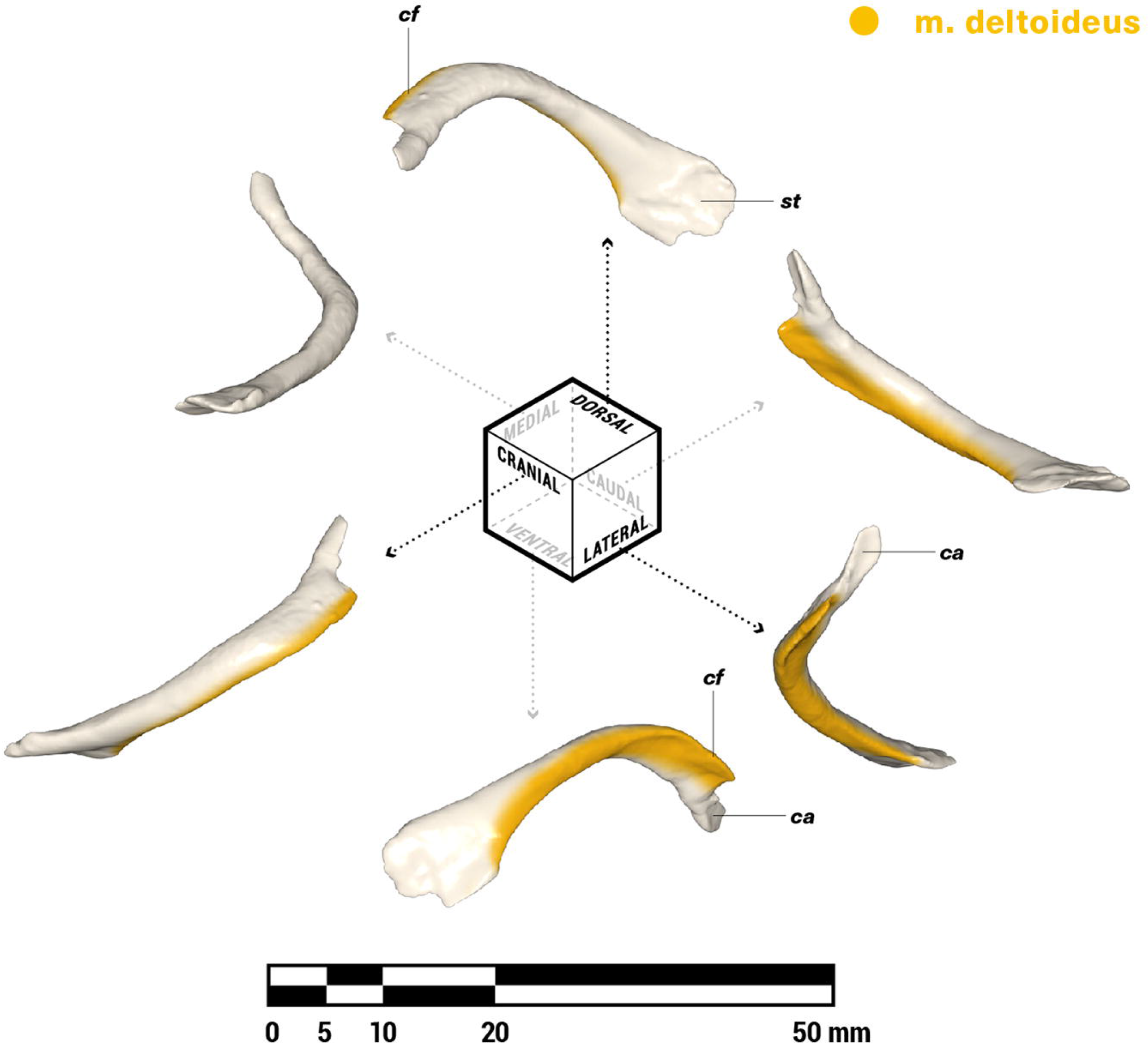
Orthographic views of the left clavicle of *M. pascuali*, with reconstructed muscle origins/insertions. Abbreviations: ca, concavity for articulation with acromion; cf, clavicular flange; st, rugose striations. Reconstructed muscles are listed in legend.

**Figure 7.**
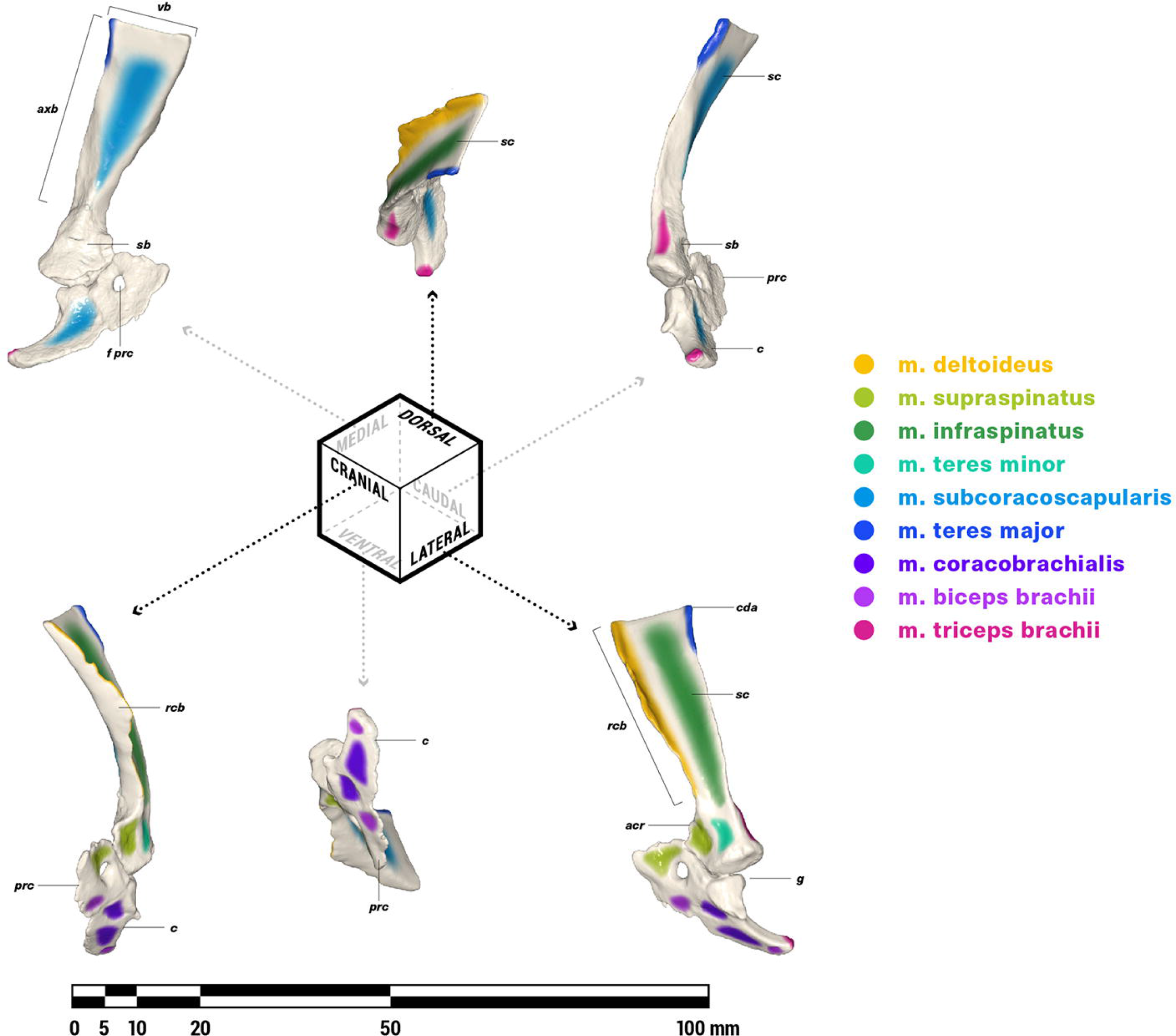
Orthographic views of the left scapulocoracoid of *M. pascuali*, with reconstructed muscle origins/insertions. Abbreviations: acr, acromion; axb, axillary border of scapula; c, coracoid (=metacoracoid *sensu* Vickaryous & Hall, 2006); cda, caudal angle of scapula; prc, procoracoid foramen; g, glenoid fossa; prc, procoracoid; rcb, reflected cranial border of scapula; sb, scapular base; sc, scapula; vb, vertebral border of scapula. Reconstructed muscles are listed in legend. Muscles are color-coded for visual differentiation, not homology.

M. deltoideus clavicularis (*sensu* Diogo et al. 2009) originates on the ventral half of the cranial border of the scapula surrounding the acromion in crocodylians (Meers, 2003), on the interclavicle extending onto the clavicle in lepidosaurs (Romer, 1922; Jenkins & Goslow, 1983) and monotremes (Howell, 1937a; Gambaryan et al. 2015), and solely along the length of the clavicle in therians (Parsons, 1896; Stein, 1981; Jenkins & Weijs,1979). The cranial edge of the clavicle in *Massetognathus* forms a distinct ridge, which extends into a protruding, anteriorly-directed flange along the distal half of the bone (cf, Fig. 6). This is the likely origin of m. deltoideus clavicularis, though it may also extend ventrally beyond the clavicle to the area of the scapula surrounding the acromion (not reconstructed). M. deltoideus clavicularis inserts with m. deltoideus scapularis, on the anterodorsal surface of the humeral deltopectoral crest (Fig. 4).

### M. supraspinatus and m. infraspinatus (Figs. 4, 7)

There is some question as to whether m. supraspinatus and m. infraspinatus were present as separate, differentiated muscles in non-mammalian cynodonts, though ontogeny shows that both are likely derivatives of the m. supracoracoideus present in non-mammals (Cheng, 1955; Romer, 1956). Some workers regard the majority of the lateral surface of the cynodont scapula as an infraspinous fossa for the origin of m. infraspinatus, with m. supraspinatus occupying the area at the craniolateral base of the scapula and the caudodorsal half of the procoracoid, where the ancestral supraspinatus attached (Gregory & Camp, 1918; Romer, 1922; Jenkins, 1971a). On the other hand, Kemp (1980a, 1980b) considered the “infraspinous fossa” an attachment site for m. deltoideus scapularis and m. teres minor. Under this hypothesis, the ventral procoracoid area attributed to m. supraspinatus by others would instead be occupied by an undifferentiated m. supracoracoideus. This latter interpretation more closely resembles the monotreme condition, wherein m. supraspinatus and m. infraspinatus are located at the cranial base of the scapula and on the procoracoid (Howell, 1937a; Gambaryan et al. 2015). However, despite their more stem-ward position in the mammal phylogeny, the suitability of extant monotremes as cynodont analogues may be compromised by modifications for a fossorial or aquatic lifestyle (Howell, 1937b; Jenkins, 1971a; Kemp, 1980b). The probable cranial migration of m. deltoideus scapularis in cynodonts (see above) likely corresponded to a dorsal expansion of m. supracoracoideus along the large, laterally-facing surface of the scapula, paralleling its origin from the lateral scapular base and procoracoid of lepidosaurs and archosaurs (Table 2).

Kemp (1980a) further argued that m. supraspinatus preceded m. infraspinatus in differentiating from m. supracoracoideus, via dorsal migration onto the anteriorly-facing surface of the reflected cranial scapular border. The ventral border of the clavicle is closely juxtaposed with the dorsal border of the procoracoid in the neutrally-posed pectoral girdle of *Massetognathus* (Fig. 3), leaving little space in between to accommodate such a muscle or its tendon, which in any case would have had to wrap around the acromion to reach Kemp’s proposed insertion on the greater tubercle of the humerus. The anteriorly-facing surface of the reflected cranial border was more likely occupied by various extrinsic muscles inserting on the scapula, such as m. trapezius and m. levator scapulae, both of which are likely plesiomorphic for amniotes (Diogo et al. 2009; Jouffroy et al. 1971).

We follow Romer (1922), Gregory and Camp (1918), and Jenkins (1971a) in reconstructing m. infraspinatus on most of the lateral surface of the scapula, caudal to the origin of m. deltoideus scapularis (Fig. 7). This muscle has a tendinous insertion on a rugosity on the distal portion of the greater tubercle, between the insertion of m. supraspinatus and the proximalmost margin of the humeral deltopectoral crest (Fig. 4). This position is intermediate between the insertion of m. supracoracoideus in *Varanus* (Jenkins & Goslow, 1983) on the proximal border of the deltopectoral crest, and the insertion of mm. spinati in mammals on the greater tubercle proper (Leach, 1977; Warburton et al. 2014; Jenkins & Weijs, 1979). An m. supracoracoideus/m. supraspinatus was likely present in *Massetognathus*, originating on the rugose area around the cranial scapular base and the adjoining procoracoid (Fig. 7). The presumptive m. supraspinatus inserts by a tendon on a rugosity on the proximal half of the humeral greater tubercle, just proximal to the insertion of m. infraspinatus (Fig. 4).

### M. teres minor (Figs. 4, 7)

According to Diogo et al. (2009), the origins of the mammalian m. teres minor are murky, with workers proposing homology with either m. scapulohumeralis anterior based on development (Romer, 1944; Cheng, 1955), or m. deltoideus scapularis based on the co-existence of m. teres minor with m. scapulohumeralis anterior in extant monotremes (Howell, 1937a; Jouffroy et al. 1971). Presuming homology with m. scapulohumeralis anterior, Romer (1922) reconstructed the origin of m. teres minor in cynodonts at the caudolateral base of the scapula, just cranial to the origin of m. triceps brachii. Gregory and Camp (1918) and Jenkins (1971a) also placed m. teres minor at the caudolateral base of the scapula, but ventral to the origin of m. triceps brachii rather than adjacent to it. Following the deltoid origin hypothesis for m. teres minor, Kemp (1980b) favored an origin high up near the vertebral border on the lateral surface of the scapula, recalling this muscle’s location in monotremes (Howell, 1937a; Gambaryan, 2015). *Massetognathus* presents no clear area of origin for m. teres minor on the dorsolateral surface of the scapula, but does possess a scar at the base of the lateral scapular surface (Fig. 7). We therefore agree with Gregory and Camp (1918) and Romer (1922) that this was the likely site of origin for m. teres minor. It is worth noting that this location is compatible with both hypotheses of origin, via either ventral differentiation of the deltoid complex or direct homology with m. scapulohumeralis anterior. In mammals, m. teres minor inserts via a tendon on the greater tubercle of the humerus (Howell, 1937a; Gambaryan et al. 2015; Jenkins & Weijs 1979; Stein, 1981; Leach, 1977). In extant lepidosaurs, m. scapulohumeralis anterior inserts on the dorsal surface of the humerus near the insertions of m. deltoideus and m. latissimus dorsi (Romer, 1922; Miner, 1925; Holmes, 1977; Jenkins & Goslow, 1983). We follow Jenkins (1971a) and Kemp (1980b) in placing the insertion of m. teres minor on a short ridge extending parallel to the long axis of the humerus from the junction of the deltopectoral crest and the greater tubercle (Fig. 4).

### M. subcoracoscapularis/subscapularis (Figs. 4, 7)

All tetrapods possess either m. subcoracoscapularis or m. subscapularis in the form of a muscle originating over much of the medial surface of the scapulocoracoid or scapula (Table 2). In lepidosaurs and monotremes, this muscle has an additional head originating on the medial surfaces of the coracoid and the procoracoid (Jenkins, 1971a; Jenkins & Goslow, 1983; Gambaryan et al. 2015). Regardless of origin, m. subcoracoscapularis/subscapularis always inserts via a tendon in the vicinity of the humeral lesser tubercle (Table 2). M. subcoracoscapularis is found on the medial side of the scapulocoracoid in all cases except for monotremes, where the subscapular fossa has migrated around the caudal border of the scapula to face posterolaterally, exposing the subscapularis in lateral view. The cynodont scapula exhibits no such torsion, and the presumptive fossa for m. subcoracoscapularis faces primarily medially, as is the case for all other tetrapods. We follow Gregory and Camp (1918), Jenkins (1971a) and Kemp (1980a) in reconstructing a two-headed subcoracoscapularis originating on the medial surfaces of the scapula and coracoid (Fig. 7), and inserting via a tendon on a rugose area at the apex of the lesser tubercle on the humerus (Fig. 4).

### M. teres major (Figs. 4, 7)

M. teres major (Table 2) is present in crocodylians (Meers, 2003) and all mammals (Howell, 1937a; Gambaryan et al. 2015; Jenkins & Weijs, 1979; Stein, 1981, 1986; Leach, 1977; George, 1977; Abdala & Diogo, 2010), but is absent in lepidosaurs (Romer, 1944; Diogo et al. 2009; Abdala & Diogo, 2010). Abdala and Diogo (2010) considered m. teres major a derivative of m. subcoracoscapularis, homologous across crocodylians and mammals, and secondarily lost in lepidosaurs and bird-line archosaurs. In therians and crocodylians, the origin of m. teres major runs dorsoventrally along the axillary border of the scapula from the caudal angle (Fig. 7), or on the lateral surface of the scapula adjacent to the axillary border (Meers, 2003; Howell, 1937a; Gambaryan et al. 2015; Jenkins & Weijs, 1979; Stein, 1981; Leach, 1977; George, 1977; Abdala & Diogo, 2010; Harvey & Warburton, 2010; Taylor, 1978). In monotremes, the origin of m. teres major runs craniocaudally along the lateral surface of the scapula, terminating at the caudal angle. Depending on whether the crocodylian m. teres major is homologous to that of mammals, m. teres major is either a level I or a level II inference. In certain cynodonts, such as *Cynognathus*, part of the axillary border of the scapula is reflected laterally into a ridge dividing the caudalmost part of the lateral scapular surface from the infraspinous fossa (Romer, 1922; Jenkins, 1971a; Liu, 2007). Some workers have interpreted this clearly demarcated fossa as the origin of m. teres major (Gregory & Camp, 1918; Jenkins, 1971a). In other cynodonts, including cynognathians such as *Luangwa* (Kemp, 1980b) and *Massetognathus*, this caudal fossa is absent, although the scapula does have a somewhat thickened area on the laterally-reflected axillary border (Fig. 7). We follow Kemp (1980b) in reconstructing m. teres major as a straplike muscle originating as a narrow strip along this thickened dorsal region of the scapula’s axillary border (Fig. 7). M. teres major likely inserted along a ridge running proximodistally along the dorsal surface of the humeral diaphysis (Fig. 4), parallel to the insertion of m. latissimus dorsi but slightly proximal (Jenkins, 1971a).

### M. coracobrachialis (Figs. 4, 7)

M. coracobrachialis (Table 2) is present in all extant tetrapods as a muscle running from the posterior part of the lateral coracoid surface—with a second head at the craniolateral base of the scapula in crocodylians (Meers, 2003)—to an insertion on the ventromedial surface of the humerus, extending onto the posteromedial surface of the humeral deltopectoral crest (Walthall & Ashley-Ross, 2006; Meers, 2003; Diogo et al. 2009; Abdala & Diogo, 2010; Holmes, 1977; Miner, 1925; Howell, 1937a; Gambaryan et al. 2015). While the presence of m. coracobrachialis as a whole is conserved among tetrapods, its subdivisions and by extension its distal attachments are not. This muscle exists as longus, medius, and brevis divisions in most amphibians (but not all, see Walthall & Ashley-Ross, 2006 and Abdala & Diogo, 2010). Lepidosaurs have lost the medius division (Abdala & Diogo, 2010, Jenkins & Goslow, 1983); crocodylians have lost all but the brevis division (Meers, 2003); monotremes seem to have lost either the brevis division (Diogo et al. 2009) or the medius (Gambaryan et al. 2015); while therians lose the longus and sometimes also the brevis (Diogo et al. 2009; Leach, 1977; George, 1977; Harvey & Warburton, 2010). While the homology of the various m. coracobrachialis divisions among extant tetrapods is beyond the scope of this paper, it seems safe to say that non-mammalian cynodonts probably had some form of m. coracobrachialis. Here we have reconstructed two origins and two insertions, representing possible brevis/medius and longus divisions.

Romer (1922) and Jenkins (1971a) considered the cynodont m. coracobrachialis to originate just caudal to m. biceps brachii within a fossa on the lateral surface of the coracoid, while Gregory and Camp (1918) assigned that fossa to m. biceps brachii and placed m. coracobrachialis on the caudal tip of the coracoid instead. *Massetognathus* has a well-marked fossa on the coracoid immediately cranial and inferior to the glenoid, and a smaller, shallower scar on the procoracoid immediately cranial to the procoracoid-coracoid suture (Fig. 7). It seems likely that m. coracobrachialis originated on the former and m. biceps brachii on the latter, echoing the arrangement of these muscles in extant *Iguana* (Romer, 1922) and *Alligator* (Meers, 2003). A second m. coracobrachialis head may have originated on the lateral surface of the coracoid, caudal and inferior to the glenoid (Fig. 7). There is no rugosity associated with m. coracobrachialis medius on the humerus of *Massetognathus*, but insertion can reasonably be assumed to have occurred on the large fossa on the ventromedial surface, with a possible insertion for m. coracobrachialis longus occurring further distal on a ridge near the entepicondyle (Fig. 4), consistent with past reconstructions (Watson, 1917; Gregory & Camp, 1918; Romer, 1922; Miner, 1925).

### M. biceps brachii (Figs. 7, 8)

M. biceps brachii is likely an amniote synapomorphy, derived from m. coracobrachialis (Abdala & Diogo, 2010). The two heads of this muscle generally originate on adjacent areas of the lateral coracoid (Miner, 1925; Romer, 1922; Jenkins, 1971a; Howell, 1937a; Holmes, 1977), although m. biceps brachii brevis is usually absent in *Alligator* (Meers, 2003), and the origin of m. biceps brachii brevis is shifted caudally to the tip of the coracoid in *Ornithorhynchus*, similar to the condition seen in *Tupaia* (George, 1977) and *Homo* (Netter et al. 1989). We follow Romer (1922) and Jenkins (1971a) in reconstructing an origin for m. biceps brachii in a depression on the lateral surface of the procoracoid, cranial to the procoracoid-coracoid suture and inferior to the procoracoid foramen (Fig. 7). A second head (m. biceps brachii brevis) may have originated on a scar on the lateral surface of the coracoid tip (Kemp, 1980b). M. biceps brachii inserts via a tendon on or near the radial tuberosity of the radius in all tetrapods (Table 2), and is reconstructed similarly in *Massetognathus* (Fig. 4).

### M. triceps brachii (Figs. 4, 7, 8)

M. triceps brachii is present in all tetrapods (Table 2). Despite the name, triceps divisions vary in number from four in urodeles, lepidosaurs, and mammals (if m. dorsoepitrochlearis is an m. triceps derivative) to five in crocodylians (Diogo et al. 2009; Abdala & Diogo, 2010). Holmes (1977) and Abdala and Diogo (2010) considered a complement of four comprising a coracoid head, a scapular head, and two humeral heads to be plesiomorphic for amniotes. In extant *Iguana*, m. triceps brachii coracoideus originates from a scar on the medial side of the coracoid, close to its caudal tip, while m. triceps brachii scapularis takes origin on a scar near the caudal base of the scapula (Romer, 1922). In extant monotremes, m. triceps coracoideus is absent whereas m. triceps brachii scapularis originates along a ridge dividing the infraspinous fossa from the subscapular fossa, a feature likely homologous with the axillary border of the scapula in other tetrapods given the relocation of m. subscapularis from the medial surface of the scapula to the posterolateral border (Gambaryan et al. 2015). *Massetognathus* has a scar on the caudomedial surface of the scapula just superior to the supraglenoid buttress, and another on the medial edge of the caudal tip of the coracoid. We follow Jenkins (1971a) in reconstructing an m. triceps brachii scapularis on the former, and a possible m. triceps brachii coracoideus on the latter (Fig. 7). There is reason to be skeptical about the presence of m. triceps brachii coracoideus in cynodonts: Romer (1922) also favored the supraglenoid scar as the origin of the scapular head of m. triceps brachii, but considered the coracoid head to have been lost, while Kemp (1980a) reasoned that a coracoid head for m. triceps brachialis is incompatible with an extended, horizontal humerus, and placed m. biceps brachii at the tip of the coracoid instead. There are no unambiguous sites of origin for the two humeral heads of m. triceps brachii on the humerus of *Massetognathus*, but these are likely to have originated somewhere along the medial and lateral surfaces of the humeral diaphysis as in the case of all tetrapods possessing them (Table 2), distal to the insertions of m. teres major and m. teres minor (Romer, 1922). All divisions of m. triceps brachii insert via a common tendon on the olecranon process of the ulna (Fig. 8).

**Figure 8.**
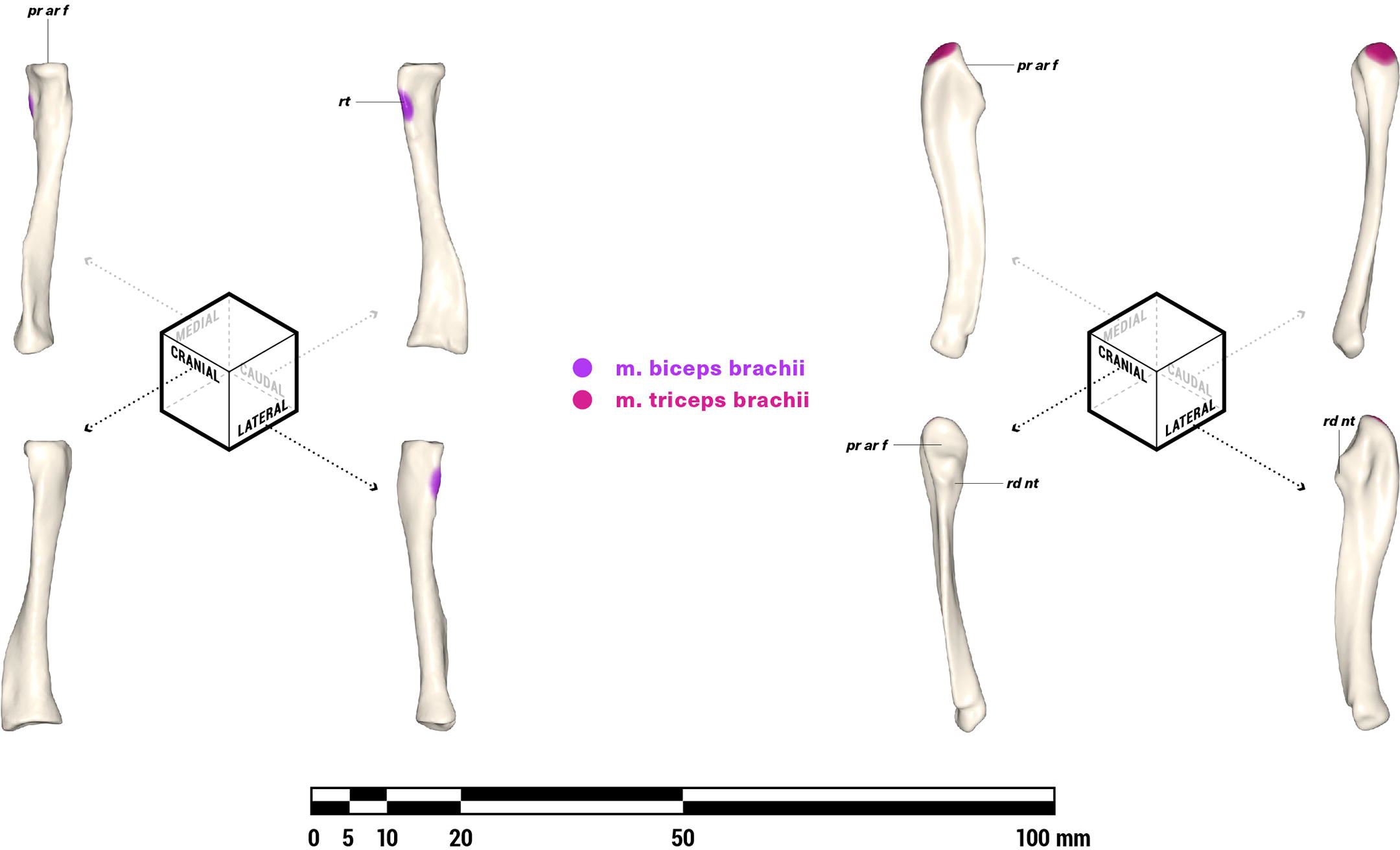
Orthographic views of the left radius and ulna of *M. pascuali*, with reconstructed muscle origins/insertions. Note that the orientations for the radius are slightly rotated from Jenkins (1971a). Abbreviations: pr ar f, proximal articular facet; rt, radial tuberosity; rd nt, radial notch. Reconstructed muscles are listed in legend.

## DISCUSSION

The present study integrates evidence from an extant phylogenetic bracket with direct observation of bony features, and corroborates earlier work (Jenkins, 1971a; Gregory & Camp, 1918) in recovering a near-therian complement of shoulder-actuating muscles in cynodonts. All but five muscles were reconstructed in *Massetognathus pascuali* as strong level I inferences, with the exceptions being m. pectoralis minor, m. deltoideus acromialis, m. teres minor, m. supraspinatus, and m. teres major. The former three are level II inferences, while the latter two are either level I or level II, depending on homology. Of these five muscles, we opted to reconstruct those whose attachments indicate distinct actions on motions of the forelimb at the shoulder (m. teres minor, m. supraspinatus, m. teres major), and omit those with similar actions at the shoulder to muscles already reconstructed as definitely present (m. pectoralis minor≈m. pectoralis major; m. deltoideus acromialis≈m. deltoideus scapularis). Our reconstructed attachment areas encompass both the excluded potential muscles and their larger neighbors, so that these muscles may be considered together as functional groups in future biomechanical analyses. Studies of extant amniotes have shown that osteological correlates to muscle attachments differ between mammalian and non-mammalian taxa (Holmes, 1977; McGowan, 1986), being more likely to manifest as rugosities in the former, versus depressions and processes in the latter (Bryant & Seymour, 1990). *Massetognathus* exhibits a combination of well-marked depressions (e.g. m. biceps brachii and m. coracobrachialis origins) and rugosities (e.g. rotator cuff and m. latissimus dorsi insertions), while distinct processes seem to be rare (coracoid origin of m. triceps brachii, if present). A mix of mammal-like and non-mammal-like muscle scars is consistent with the intermediate phylogenetic position of non-mammalian cynodonts. Notably, several of the muscles reconstructed (namely, m. pectoralis, m. deltoideus, m. latissimus dorsi, m. teres major, and m. teres minor) have long, narrow insertions extending proximodistally along the humeral diaphysis, raising the question of whether resistance to torsion in long, flat muscles might present constraints on humeral movement.

### Comparison with previous range of motion measurements

The mobility of individual joints has not been extensively documented for the cynodont forelimb, and most workers have focused on the gleno-humeral joint over more proximal or distal articulations. Jenkins (1971a) reported 30° of long-axis rotation and 40° of adduction at the gleno-humeral joint, while Kemp (1980a, 1980b) reported 90° of long-axis rotation, nearly 90° of protraction-retraction (“from almost transverse to fairly close to posteriorly directed”), and a “reasonable degree” of abduction-adduction at the same joint. Both Jenkins’ and Kemp’s numbers fall close to the limits reported here, with the exception of Kemp’s long-axis rotation measurement, which exceeds ours by 20°. The discrepancy between Jenkins’ and Kemp’s measurements may be phylogenetic, but may also reflect differing assumptions of joint kinematics. While both workers hypothesized a translational component of humeral motion, Jenkins’ values resemble our rotation-only measurements, while Kemp’s estimates are closer to our combined translation-rotation measurements (Table 1). Oliveira and Schultz (2016) measured 70° of abduction-adduction, 15-20° of retraction-protraction, and an unspecified amount of long-axis rotation at the gleno-humeral joint. Their measurements fall within our maximum ranges, although they do not report their joint space assumptions and coordinate systems.

Kemp (1980b) reported over 90° of flexion-extension for the radius and ulna at their respective articulations with the humerus. Oliveira and Schultz (2016) considered the radius and ulna as a functional unit, measuring over 100° of flexion-extension at the elbow. Oliveira and Schultz additionally reported 25° of mediolateral rotation at the clavo-interclavicular joint, and 15° of roll, 25° of yaw, and 30° of pitch at the acromio-clavicular joint. Again, these values all fall well within our measured ranges (Table 1).

It should be stressed that the angular ranges reported in Table 1 represent maximum estimates of joint mobility. It is well established that extrinsic soft tissues such as ligaments, joint capsules, labra, muscles, and integument restrict range of motion in an intact animal to a subset of the mobility assessed from manipulation of dry bones (Pierce et al. 2012; Arnold et al. 2014; Hutson & Hutson, 2012, 2014); although the shoulder appears to be less constrained than the hip, and long-axis rotation seems to be the most affected (Pierce et al. 2012). The aim of this analysis was to establish reasonable maximum ranges as a basis for future validation and refinement, and we fully expect the maximum range of motion at all joints modeled here to decrease substantially with the imposition of soft tissue constraints. Radiographic studies of *in vivo* joint utilization (e.g. Fischer, 1994; Kambic et al. 2014) suggest that an even smaller fraction of that mobility is actually employed during normal locomotion, with the remaining available joint surface reserved for non-locomotor behaviors.

### Comparison with previous muscle reconstructions

Using mammalian muscle anatomy as reference, Oliveira and Schultz (2016) reconstructed the pectoral girdle and forelimb musculature of *Trucidocynodon riograndensis*, a Brazilian Triassic eucynodont. The present reconstruction agrees with theirs in the location of the scapular and humeral heads of m. triceps brachii, and differs in the relative arrangement of m. latissimus dorsi and m. teres major insertions (m. latissimus dorsi inserts distal and medial to m. teres major in our reconstruction, whereas m. latissimus dorsi is proximal and lateral in Oliveira and Schultz, 2016), and the presence of a humeral origin for m. biceps brachii (absent in ours, present in theirs). We attribute these discrepancies to our use of extant phylogenetic bracketing for determining muscle attachments, in contrast to their adherence to mammalian anatomy. Unlike Oliveira and Schultz, we stopped short of recreating the morphology of the muscles themselves. While Lautenschlager (2013) and others (Holliday, 2009; Cuff & Rayfield, 2015) have shown the feasibility of using topography and spatial exclusion to establish the morphology of tightly juxtaposed cranial muscles with direct attachments, limb muscles tend to be more widely spaced, and Bryant and Seymour (1990) caution that architecture and non-uniform cross-sections can confound three-dimensional reconstructions of muscles with tendinous attachments.

While Gregory and Camp (1918) recovered a similar muscle reconstruction to ours for the cynodont *Cynognathus*, their skeletal reconstruction differed in one important respect. *Cynognathus* was reconstructed with the scapulocoracoids much closer to the animal’s sagittal plane, such that the coracoids appear to contact the interclavicle along the ventral midline. This arrangement resembles that seen in extant monotremes, wherein the coracoids articulate with the interclavicle and the procoracoids are closely apposed, occasionally overlapping asymmetrically (Cave, 1970). This has been suggested to be a derived condition allowing better resistance to compressive forces, and possibly related to fossorial or swimming behaviors (Luo, 2015). In *Massetognathus*, the length and curvature of the clavicles necessitate substantial separation between the scapulocoracoids, regardless of clavicular mobility. The lateral separation between the scapulocoracoids and the interclavicle tends to get understated in two-dimensional reconstructions, many of which depict a lateral view showing only the smaller vertical component of the gap (e.g. Jenkins, 1971a; Kemp, 1980a, 1980b; Sun & Li, 1985).

Gregory and Camp (1918) noted that the suprascapular cartilages in their reconstruction are probably too small, as the dorsalmost extent of these structures is still far ventral to the tops of the neural spines in the vertebral column, possibly compromising the ability of m. rhomboideus and m. trapezius to suspend the thorax. Greater separation between the scapulocoracoids would ameliorate this by placing the scapulae higher up on the animal’s body wall. Curiously, Gregory and Camp go on to hypothesize a thin epicoracoid element in the Permian therapsid *Moschops*, spanning the gap between the clavicles, procoracoids, and the interclavicle. It is unclear why they did not propose a similar structure in *Cynognathus*, but the irregular ventromedial margins of the procoracoid and coracoid (Fig. 7) in *Massetognathus* are consistent with having possibly articulated with unossified epicoracoid cartilages in life, as seen in extant lepidosaurs (Fürbringer, 1900). The wide separation between the scapulocoracoids and the midventral interclavicle in *Massetognathus* is suggestive of a functional transformation away from the massive, heavily ossified, “U”-shaped girdles seen at the base of the synapsid tree, possibly reflecting a shift in the loading regime experienced by the forelimb and pectoral girdle from mediolateral compression towards more vertically-oriented reaction forces (Jenkins, 1971a). Continued reduction of the endochondral primary girdle (scapulocoracoid) and dermal secondary girdle (clavicle, interclavicle) throughout cynodont evolution may have resulted in progressively greater separation between the scapulocoracoids, presaging the fully independent therian pectoral girdle, dominated by a large scapula and a reduced, strutlike clavicle (Jenkins & Weijs, 1979; Luo, 2015).

### Mobility of the pectoral girdle

Kemp (1980b) hypothesized that some degree of mobility in the cynodont pectoral girdle would have been necessary in order to increase the functional length of the forelimb to match that of the hindlimb. Hopson (2015) also considered the issue of stride length in his analysis of the pelycosaur *Dimetrodon*, attributing it to mediolateral rotation of the pectoral girdle from increased bending of the trunk. Lateral bending was suggested to have been lost or greatly reduced in therapsids (Jenkins, 1971a; Kemp, 2005), and it seems unlikely that the cynodont pectoral girdle could have rotated as a single unit with respect to the axial skeleton. It is worth noting that multiple derived amniote lineages seem to have converged on some degree of pectoral girdle mobility—substantial translational coracosternal mobility has been reported in several extant non-mammals, including lepidosaurs (Peterson, 1973; Jenkins & Goslow, 1983), ornithodirans (Baier et al. 2013), and pseudosuchians (Baier & Gatesy, 2013). Little is yet known about the soft-tissue constraints acting on the joints of the non-mammalian synapsid pectoral girdle. Based on osteology, there is no *a priori* reason to reject the possibility that rotation at the clavo-interclavicular and acromio-clavicular joints may have furnished a degree of independent scapulocoracoid mobility in *Massetognathus*. Experimenting with hypothetical, non-physiological poses reveals that scapulocoracoid mobility allows a significant increase in maximum forelimb excursion, as well as slightly more medial placement of the wrist (Fig. 9). Among derived probainognathian cynodonts, tritylodonts (though not tritheledonts), basal mammaliaforms, and monotremes exhibit expanded lateral processes on their interclavicles that overlap substantially with the proximal ends of their clavicles, precluding clavo-interclavicular mobility (Luo, 2015; Jenkins & Parrington, 1976). The ability to unilaterally move one side of the pectoral girdle may have helped functionally separate limb pairs, possibly laying the groundwork for the evolution of asymmetrical gaits. However, if clavo-interclavicular mobility was indeed present in more stem-ward cynodonts, the robust, immobile pectoral girdle of monotremes would then represent an interesting atavistic reversal (Ji et al. 1999).

**Figure 9.**
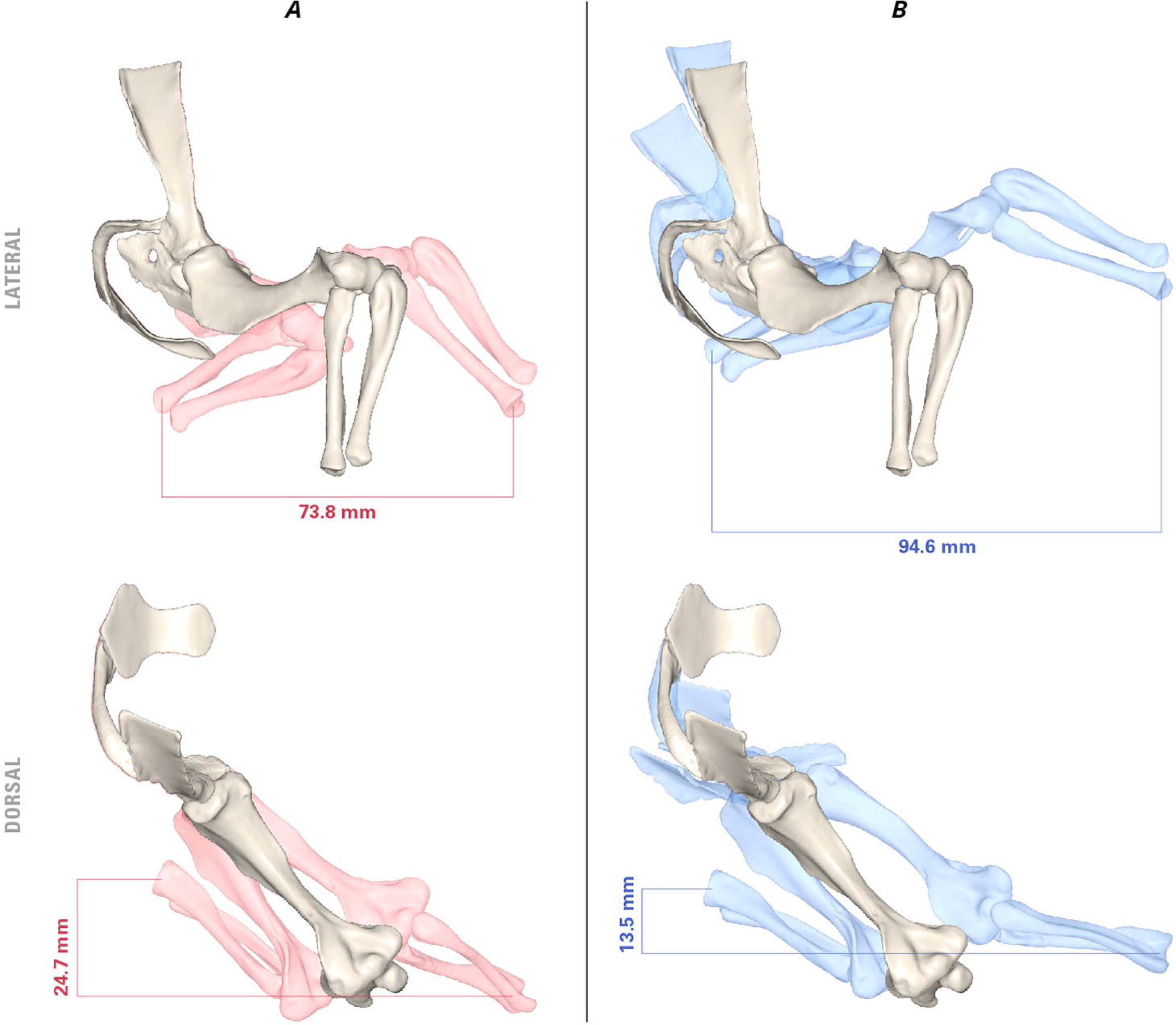
Potential pectoral limb excursion with fixed (A) and mobile (B) clavo-interclavicular and acromio-clavicular articulations. Neutral pose in white. Joint angles (protracted): [clavicle rotated medially 15°; scapulocoracoid yawed laterally 15°, rolled laterally 5°, pitched caudally 5°]; humerus adducted 20°, supinated 15°, protracted 15°; radius and ulna flexed 45°. Joint angles (retracted): [clavicle rotated laterally 15°; scapulocoracoid rolled medially 5°, pitched cranially 30°]; humerus pronated 10°, retracted 15°; radius and ulna extended 45°.

### Epiphyses and range of motion

Among amniotes, ossified epiphyses with well-defined articular surfaces arose convergently in mammals and lepidosaurs (Haines, 1969); unossified epiphyses lacking growth plates are plesiomorphic for cynodonts (Luo et al. 2007b), and the articular surfaces of long bones are thought to have been extended and elaborated by cartilaginous structures in life (Jenkins, 1971a; Jenkins & Parrington, 1976). Progressive removal of connective tissues in archosaurs has shown that cartilage may increase the range of motion available at a joint (Hutson & Hutson, 2012), and that the surface morphology of cartilage caps may differ sufficiently from that of the underlying bone to meaningfully alter joint action (Holliday et al. 2010). While it is unclear how cynodont cartilage caps compared to those of either extant crocodylians or extant avians in thickness, both Jenkins and Kemp mentioned them in their reconstructions of cynodont limb function. Jenkins (1971) noted the discrepancy in curvature between the “notch-shaped” hemisellar glenoid and the convex, ovoid humeral head, and posited that a substantial thickness of cartilage must have been present in life to increase congruence between the articular surfaces, much like in extant archosaurs (Holliday et al. 2010). Kemp (1980b), on the other hand, argued that any cartilaginous intermediary would have had to be “absurdly thick” to match the humeral head’s curvature to that of the glenoid fossa, and interpreted the incongruity in articular morphology as representing a “rolling” rather than sliding articulation at the gleno-humeral joint.

The 0.25 mm of epiphyseal cartilage modeled here (both proximal and distal) is based on mammalian measurements from the literature. An articular cap measuring 0.25 mm on each epiphysis adds up to 0.5 mm per long bone, and represents an approximate 1% contribution to humeral length in *Massetognathus*. This is comparable with measurements taken by Holliday et al. (2010) from extant avians (0.07%-3.72%), but falls below the 7.99% measured in *Alligator*. However, it is unclear whether archosaurs, and *Alligator* in particular, represent good models for reconstructing non-mammalian synapsids. We were unable to locate cartilage measurements for lepidosaurs or amphibians in the literature, but epiphyseal cartilage is likely to be thin in the former clade given the presence of extensive secondary ossification centers (Haines, 1969), and extensive in the latter due to their aquatic tendencies.

Greater cartilage thickness—and consequently greater joint space—may indeed further increase range of motion. An informal sensitivity test on this model revealed that a threefold increase in joint space (from 0.50 mm to 1.50 mm) increased rotation-only range of motion to ranges comparable to combined translation-rotation levels measured with the original 0.50 mm joint space. However, thicker cartilage may significantly alter functional articular morphology from that of the underlying bone (Holliday et al. 2010). Since the superficial cartilage morphology is unpreserved, we are unable to satisfactorily bound this into a testable problem, and instead present measurements based on a joint space of 0.5 mm (Table 1) a conservative estimate.

### Evolution toward the therian glenoid fossa

The separate scapular and coracoid facets that together comprise the glenoid fossa deserve closer examination. The basal cynodont glenoid is commonly described as laterally or posterolaterally-facing and either “hemisellar” (Jenkins, 1971a) or “notch”-shaped (Kemp, 2005), a morphology retained through progressively more derived eucynodonts and mammaliaforms before finally evolving into the familiar, ventrally-directed, mammalian ball-and-socket joint in the Jurassic theriimorphs (Luo, 2015). In *Massetognathus*, the scapular and coracoid facets of the glenoid are dissimilar in morphology (Fig. 10). Viewed together, the two create the impression of a unified notch, but close examination reveals that the coracoid facet is mediolaterally convex, with a small, laterally projecting lip at its ventral apex creating a small degree of dorsoventral concavity. The scapular facet is set at an obtuse angle (approx. 130°) to the coracoid facet, and is slightly concave at the center (*contra* Jenkins, 1970b, where it is described as slightly convex in the same specimen). A procoracoid contribution to the glenoid seems somewhat variable among cynognathians; it is completely excluded from the glenoid in *Massetognathus*, but makes up a small, cranial portion of the articulation in *Cynognathus* (Jenkins, 1971a).

**Figure 10.**
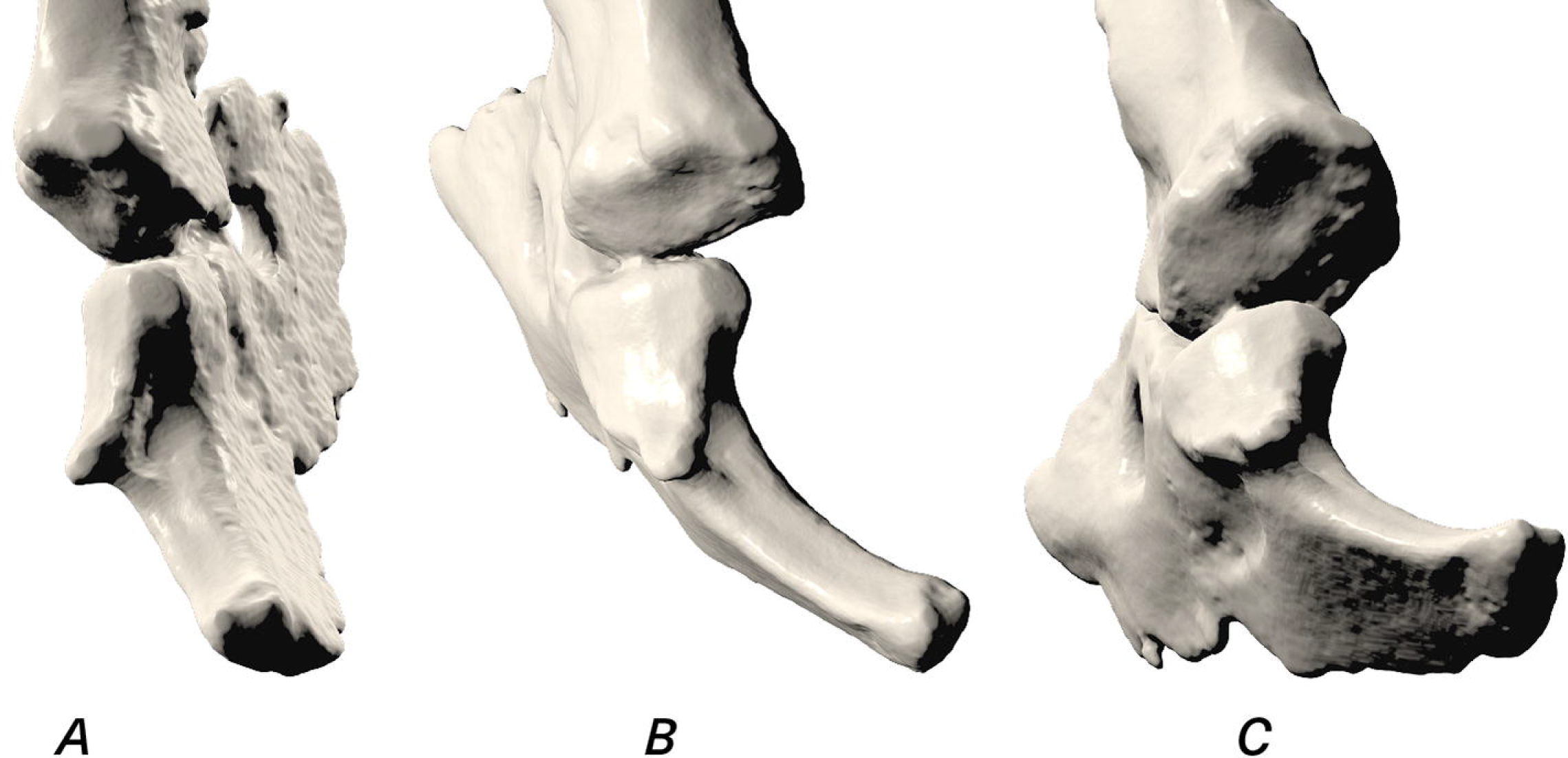
Close-up caudal (A), caudolateral (B) and ventrocaudolateral (C) views of the glenoid fossa of *M. pascuali*, showing the convex coracoid facet and the concave scapular facet.

Jenkins (1971a) interpreted the separate scapular and coracoid facets as representing a functional division within the glenoid, with a possibly ventrally-facing scapular facet serving to transmit the vertical component of ground reaction force during locomotion. The full-girdle reconstruction of *Massetognathus* presented here confirms a ventral orientation for the scapular facet (Fig. 3). Taking Jenkins’ logic a step further, one could hypothesize that the laterally-facing coracoid and procoracoid may have initially functioned to accommodate a substantial lateral component of ground reaction force. Increasingly parasagittal limb kinematics and increasingly vertically-oriented ground reaction forces in more crown-ward taxa might have accompanied progressively reduced coracoid and procoracoid contributions to the glenoid, until all that remained was a broadened, deepened scapular facet forming a ventrally-facing, socket-shaped glenoid. Under this scenario, the characteristic therian ball-and-socket joint was achieved not through reorientation of an ancestrally laterally-facing composite glenoid as a whole, but rather progressive reduction, ventral reorientation, and possibly partial assimilation (Vickaryous & Hall, 2006) of the plesiomorphic coracoid facet into a pre-existing, ventrally-facing, concave scapular facet. This hypothesis of glenoid transformation is wholly compatible with macroevolutionary trends in coracoid reduction and scapular expansion among derived cynodonts, as reported by Luo (2015) as well as Jenkins and Weijs (1979).

### Differences between monotreme and cynodont pectoral girdles

As the sister group to crown therians, extant monotremes have long been used as models for understanding non-mammalian cynodont biology (Gregory & Camp, 1918; Kemp, 1980a; Luo, 2007). In addition to the foregoing discrepancy in scapulocoracoid position, our reconstruction of *Massetognathus* differs from monotremes in a number of important musculoskeletal respects, namely the length of the clavicles and the extent of their overlap with the interclavicle; the significant mediolateral torsion present in the monotreme scapula but absent in *Massetognathus;* the higher aspect ratio and weaker long-axis torsion of the humerus in *Massetognathus*; the insertion of m. latissimus dorsi (much more distal in monotremes); the origin of m. teres minor (far dorsal in monotremes); and the orientation of the fossa for m. subcoracoscapularis (medially-facing in *Massetognathus* and all other amniotes, posterolaterally-facing in monotremes). In reconstructing *Massetognathus*, we found that monotreme muscle attachments were frequently the exception in the extant phylogenetic bracket, with the monotreme m. subcoracoscapularis in particular differing from its location and attachments in therians and saurians alike. These discrepancies raise the question of whether ecological specializations may have left a confounding signature on the musculoskeletal organization of the monotreme pectoral girdle and forelimb. It seems reasonable to entertain the possibility that niche adaptation and low ecological diversity in extant monotremes may render them compromised locomotor analogues for non-mammalian cynodonts (Howell, 1936; Jenkins, 1971a; Jenkins, 1989). If this is the case, it may be prudent to situate interpretations of postcranial function in non-mammalian cynodonts within a wide, comparative context spanning mammalian and non-mammalian forms.

## CONCLUSION

The musculoskeletal reconstruction of *Massetognathus pascuali* presented here recovers maximum ranges of motion and muscle origins and insertions comparable to those found in previous studies, and reveals features of the pectoral girdle not previously described for this genus, such as a wide separation between the scapulocoracoids and the interclavicle, possible mobility at the clavo-interclavicular and acromio-clavicular joints, and a ventrally-facing, concave scapular facet of the glenoid fossa. Extensive functional interpretation of our 3D reconstruction awaits further comparison with extant mammalian and non-mammalian taxa. Having already mapped well-defined muscle attachment areas onto a digital skeleton, we have laid the groundwork for constructing interactive musculoskeletal models. Future experiments that combine detailed musculoskeletal modeling with *ex vivo* and *in vivo* data from extant taxa will provide the opportunity to investigate the relationship between skeletal posture and muscle function, and shed further light on the cynodont pectoral limb and its significance to the rise of mammals.

## ACKNOWLEDGEMENTS

We thank Katrina Jones for her help with CT scanning, and Jessica Cundiff for assistance with specimens in the MCZ. The Pierce and Biewener labs provided valuable feedback. Financial support was provided by Harvard University funds made available to S.E.P. CT data is reposited in the Department of Vertebrate Paleontology, Museum of Comparative Zoology, Harvard University. The authors declare no conflicts of interest.

## AUTHOR CONTRIBUTIONS

All authors contributed to study conception and manuscript preparation. P.H.L. and S.E.P. contributed to experimental design, data acquisition, and interpretation. All authors gave final approval for publication.

---

**Supplementary Figure 1.** Three-dimensional reconstruction of the left pectoral limb of *M. pascuali*. Reconstructed muscle origins/insertions are listed in legends of Figs. 4-8. This interactive PDF may be viewed in Adobe Acrobat (Adobe Systems Incorporated, San Jose, CA, USA).

## REFERENCES

Abdala, V., & Diogo, R. (2010). Comparative anatomy, homologies and evolution of the pectoral and forelimb musculature of tetrapods with special attention to extant limbed amphibians and reptiles. Journal of Anatomy, 217(5), 536–573.

Arnold, P., Fischer, M. S., & Nyakatura, J. A. (2014). Soft tissue influence on ex vivo mobility in the hip of Iguana: comparison with in vivo movement and its bearing on joint motion of fossil sprawling tetrapods. Journal of Anatomy, 225(1), 31–41.

Baier, D. B., Gatesy, S. M., & Dial, K. P. (2013). Three-dimensional, high-resolution skeletal kinematics of the avian wing and shoulder during ascending flapping flight and uphill flap-running. PloS One, 8(5), e63982.

Baier, D. B., & Gatesy, S. M. (2013). Three-dimensional skeletal kinematics of the shoulder girdle and forelimb in walking Alligator. Journal of Anatomy, 223(5), 462–473.

Bates, K. T., & Schachner, E. R. (2012). Disparity and convergence in bipedal archosaur locomotion. Journal of the Royal Society, Interface / the Royal Society, 9(71), 1339–1353.

Brainerd, E. L., Baier, D. B., Gatesy, S. M., Hedrick, T. L., Metzger, K. A., Gilbert, S. L., & Crisco, J. J. (2010). X-ray reconstruction of moving morphology (XROMM): precision, accuracy and applications in comparative biomechanics research. Journal of Experimental Zoology. Part A, Ecological Genetics and Physiology, 313(5), 262–279.

Bryant, H. N., & Seymour, K. L. (1990). Observations and comments on the reliability of muscle reconstruction in fossil vertebrates. Journal of Morphology, 206(1), 109–117.

Cave, A. J. E. (1970). Observations on the monotreme interclavicle. Journal of Zoology, 160(3), 297–312.

Cheng, C.-C. (1955). The development of the shoulder region of the opossum, Didelphys virginiana, with special reference to the musculature. Journal of Morphology, 97(3), 415–471.

Coues, E. (1871). On the Myology of the Ornithorhynchus.

Coues, E. & Wyman, J. (1872). The osteology and myology of Didelphys virginiana (p. 116). Boston: The Society,.

Cuff, A. R., & Rayfield, E. J. (2015). Retrodeformation and muscular reconstruction of ornithomimosaurian dinosaur crania. PeerJ, 3, e1093.

Davison, A. (1895). A contribution to the anatomy and phylogeny of Amphiuma means (Gardner). Journal of Morphology, 11(2), 375–410.

Delp, S. L., & Loan, J. P. (1995). A graphics-based software system to develop and analyze models of musculoskeletal structures. Computers in Biology and Medicine, 25(1), 21–34.

Diogo, R., Abdala, V., Aziz, M. A., Lonergan, N., & Wood, B. A. (2009). From fish to modern humans--comparative anatomy, homologies and evolution of the pectoral and forelimb musculature. Journal of Anatomy, 214(5), 694–716.

Field, D. A. (1988). Laplacian smoothing and Delaunay triangulations. Communications in Applied Numerical Methods, 4(6), 709–712.

Fischer, M. S. (1994). Crouched posture and high fulcrum, a principle in the locomotion of small mammals: The example of the rock hyrax (Procavia capensis) (Mammalia: Hyracoidea). Journal of Human Evolution, 26(5–6), 501–524.

Fischer, M. S., Schilling, N., Schmidt, M., Haarhaus, D., & Witte, H. (2002). Basic limb kinematics of small therian mammals. The Journal of Experimental Biology, 205(Pt 9), 1315–1338.

Fürbringer, M. (1900). Zur vergleichenden Anatomie des Brustschulterapparates und der Schultermuskeln. IV Teil. Jena Ische Zeitschr. Naturwiss., Jena, 34, 351.

Gambaryan, P. P., Kuznetsov, A. N., Panyutina, A. A., & Gerasimov, S. V. (2015). Shoulder girdle and forelimb myology of extant Monotremata. Russian Journal of Theriology, 14(1), 1–56.

Gatesy, S. M. (1991). Hind limb movements of the American alligator (Alligator mississippiensis) and postural grades. Journal of Zoology, 224(4), 577–588.

Gatesy, S. M., Baier, D. B., Jenkins, F. A., & Dial, K. P. (2010). Scientific rotoscoping: a morphology-based method of 3-D motion analysis and visualization. Journal of Experimental Zoology. Part A, Ecological Genetics and Physiology, 313(5), 244–261.

George, R. M. (1977). The limb musculature of the Tupaiidae. Primates; Journal of Primatology, 18(1), 1–34.

Gregory, W. K., & Camp, C. L. (1918). Studies in comparative myology and osteology. American Museum of Natural History.

Grood, E. S., & Suntay, W. J. (1983). A joint coordinate system for the clinical description of three-dimensional motions: application to the knee. Journal of Biomechanical Engineering.

Haines, R. W. (1942). THE EVOLUTION OF EPIPHYSES AND OF ENDOCHONDRAL BONE. Biological Reviews of the Cambridge Philosophical Society, 17(4), 267–292.

Harvey, K. J., & Warburton, N. (2010). Forelimb musculature of kangaroos with particular emphasis on the tammar wallaby Macropus eugenii (Desmarest, 1817). Australian Mammalogy, 32(1), 1–9.

Hildebrand, M. (1989). The quadrupedal gaits of vertebrates. Bioscience, 39(11), 766–775.

Holliday, C. M. (2009). New insights into dinosaur jaw muscle anatomy. Anatomical Record, 292(9), 1246–1265.

Holliday, C. M., Ridgely, R. C., Sedlmayr, J. C., & Witmer, L. M. (2010). Cartilaginous epiphyses in extant archosaurs and their implications for reconstructing limb function in dinosaurs. PloS One, 5(9). https://doi.org/10.1371/journal.pone.0013120

Holmes, R. (1977). The osteology and musculature of the pectoral limb of small captorhinids. Journal of Morphology, 152(1), 101–140.

Hoppe, H. (2008). Poisson Surface Reconstruction and Its Applications. In Proceedings of the 2008 ACM Symposium on Solid and Physical Modeling (pp. 10–10). New York, NY, USA: ACM.

Hopson, J. A. (2015). Fossils, Trackways, and Transitions in Locomotion: A Case Study of Dimetrodon. In Great Transformations in Vertebrate Evolution.

Howell, A. B. (1936). The phylogenetic arrangement of the muscular system. The Anatomical Record.

Howell, A. B. (1937a). Morphogenesis of the Shoulder Architecture. Part V. Monotremata. The Quarterly Review of Biology, 12(2), 191–205.

Howell, A. B. (1937b). The Swimming Mechanism of the Platypus. Journal of Mammalogy, 18(2), 217–222.

Hutchinson, J. R., Anderson, F. C., Blemker, S. S., & Delp, S. L. (2005). Analysis of hindlimb muscle moment arms in Tyrannosaurus rex using a three-dimensional musculoskeletal computer model: implications for stance, gait, and speed. Paleobiology, 31(4), 676–701.

Hutchinson, J. R., Rankin, J. W., Rubenson, J., Rosenbluth, K. H., Siston, R. A., & Delp, S. L. (2015). Musculoskeletal modelling of an ostrich (Struthio camelus) pelvic limb: influence of limb orientation on muscular capacity during locomotion. *PeerJ*, *3*, e1001.

Hutson, J. D., & Hutson, K. N. (2012). A test of the validity of range of motion studies of fossil archosaur elbow mobility using repeated-measures analysis and the extant phylogenetic bracket. The Journal of Experimental Biology, 215(Pt 12), 2030–2038.

Hutson, J. D., & Hutson, K. N. (2014). A repeated measures analysis of the effects of soft tissues on wrist range of motion in the extant phylogenetic bracket of dinosaurs: Implications for the …. The Anatomical Record. Retrieved from http://onlinelibrary.wiley.com/doi/10.1002/ar.22903/full

Jenkins, F. A. (1970a). Cynodont postcranial anatomy and the “prototherian” level of mammalian organization. Evolution; International Journal of Organic Evolution, 230–252.

Jenkins, F. A. (1970b). The Chañares (Argentina) Triassic reptile fauna VII. The postcranial skeleton of the traversodontid Massetognathus pascuali (Therapsida, Cynodontia). Breviora, 352, 1–28.

Jenkins, F. A. (1989). Monotremes and the biology of Mesozoic mammals. Netherlands Journal of Zoology.

Jenkins, F. A., & Goslow, G. E. (1983). The functional anatomy of the shoulder of the savannah monitor lizard (Varanus exanthematicus). Journal of Morphology, 175(2), 195–216.

Jenkins, F. A., Jr, & Parrington, F. R. (1976). The postcranial skeletons of the Triassic mammals Eozostrodon, Megazostrodon and Erythrotherium. Philosophical Transactions of the Royal Society of London. Series B, Biological Sciences, 273(926), 387–431.

Jenkins, F. A. (1971a). The postcranial skeleton of African cynodonts: problems in the early evolution of the mammalian postcranial skeleton. Harvard MCZ.

Jenkins, F. A. (1971b). Limb posture and locomotion in the Virginia opossum (Didelphis marsupialis) and in other non-cursorial mammals. Journal of Zoology, 165(3), 303–315.

Jenkins, F. A. (1993). The evolution of the avian shoulder joint. American Journal of Science.

Jenkins, F. A., & Weijs, W. A. (1979). The functional anatomy of the shoulder in the Virginia opossum (Didelphis virginiana). Journal of Zoology, 188(3), 379–410.

Ji, Q., Luo, Z.-X., & Ji, S. A. (1999). A Chinese triconodont mammal and mosaic evolution of the mammalian skeleton. Nature, 398(6725), 326–330.

Ji, Q., Luo, Z.-X., Yuan, C.-X., & Tabrum, A. R. (2006). A swimming mammaliaform from the Middle Jurassic and ecomorphological diversification of early mammals. Science, 311(5764), 1123–1127.

Jouffroy, F. K. L., Saban, J., Souteyrand-Boulenger, R., Jouffroy, J., & Others. (1971). Mammifères: musculature des membres, musculature peaucière, musculature des monotrèmes. Arthrologie. Retrieved from http://www.sidalc.net/cgi-bin/wxis.exe/?IsisScript=FCL.xis&method=post&formato=2&cantidad=1&expresion=mfn=001693

Kambic, R. E., Roberts, T. J., & Gatesy, S. M. (2014). Long-axis rotation: a missing degree of freedom in avian bipedal locomotion. The Journal of Experimental Biology, 217(Pt 15), 2770–2782.

Kemp, T. S. (1978). Stance and gait in the hindlimb of a therocephalian mammal-like reptile. Journal of Zoology, 186(2), 143–161.

Kemp, T. S. (1980a). Aspects of the structure and functional anatomy of the Middle Triassic cynodont Luangwa. J. Zool., Lond., 191, 193–239.

Kemp, T. S. (1980b). The Primitive Cynodont Procynosuchus: Structure, Function and Evolution of the Postcranial Skeleton. Philosophical Transactions of the Royal Society of London. Series B, Biological Sciences, 288(1027), 217–258.

Kemp, T. S. (2005). The Origin and Evolution of Mammals. OUP Oxford.

Kirsch, J. A. W. (1973). Notes for the dissection of the opossum, Didelphis virginiana. Madison, WI.

Lautenschlager, S. (2013). Cranial myology and bite force performance of Erlikosaurus andrewsi: a novel approach for digital muscle reconstructions. Journal of Anatomy, 222(2), 260–272.

Leach, D. (1977). The forelimb musculature of marten (Martes americana Turton) and fishes (Martes pennanti Erxleben). Canadian Journal of Zoology, 55(1), 31–41.

Liu, J. (2007). New traversodontid materials from North Carolina, USA and the taxonomy, phylogeny of Traversodontidae (Synapsida: Cynodontia). COLUMBIA UNIVERSITY.

Liu, J., & Abdala, F. (2014). Phylogeny and Taxonomy of the Traversodontidae. In C. F. Kammerer, K. D. Angielczyk, & J. Fröbisch (Eds.), Early Evolutionary History of the Synapsida (pp. 255–279). Springer Netherlands.

Liu, J., & Olsen, P. (2010). The Phylogenetic Relationships of Eucynodontia (Amniota: Synapsida). Journal of Mammalian Evolution, 17(3), 151–176.

Luo, Z.-X. (2007). Transformation and diversification in early mammal evolution. Nature, 450(7172), 1011–1019.

Luo, Z.-X., Chen, P., Li, G., & Chen, M. (2007b). A new eutriconodont mammal and evolutionary development in early mammals. Nature, 446(7133), 288–293.

Luo, Z.-X. (2015). Origin of the Mammalian Shoulder. In Great Transformations in Vertebrate Evolution.

McGowan, C. (1986). The wing musculature of the weka. Gallirallus Australis.

Meers, M. B. (2003). Crocodylian forelimb musculature and its relevance to Archosauria. The Anatomical Record. Part A, Discoveries in Molecular, Cellular, and Evolutionary Biology, 274(2), 891–916.

Meng, J., Hu, Y., Wang, Y., Wang, X., & Li, C. (2006). A Mesozoic gliding mammal from northeastern China. Nature, 444(7121), 889–893.

Miner, R. W. (1925). THE PECTORAL LIMB OF ERYOPS AND OTHER PRIMITIVE TETRAPODS. Bulletin of the AMNH, 51(7).

Mivart, S. G. (1869). Notes on the Myology of Menopoma alleghaniense. Proceedings of the Zoological Society of London, 37(1), 254–271.

Netter, F. H., Colacino, S., & Others. (1989). Atlas of human anatomy (Vol. 11). Ciba-Geigy Summit, NJ.

Nyakatura, J. A., Allen, V. R., Lauströer, J., Andikfar, A., Danczak, M., Ullrich, H.-J., … Fischer, M. S. (2015). A Three-Dimensional Skeletal Reconstruction of the Stem Amniote Orobates pabsti (Diadectidae): Analyses of Body Mass, Centre of Mass Position, and Joint Mobility. PloS One, 10(9), e0137284.

Oliveira, T. V. D., & Schultz, C. L. (2016). Functional Morphology and Biomechanics of the Cynodont Trucidocynodon riograndensis from the Triassic of Southern Brazil: Pectoral Girdle and Forelimb. Acta Palaeontologica Polonica, 61(2), 377–386.

Parsons, F. G. (1896). 4. Myology of Rodents.—Part II. An Account of the Myology of the Myomorpha, together with a Com parison of the Muscles of the various Suborders of Rodents. In Proceedings of the Zoological Society of London (Vol. 64, pp. 159–192). Wiley Online Library.

Peterson, J. A. (1973). Adaptation for arboreal locomotion in the shoulder region of lizards. University of Chicago Press.

Pierce, S. E., Clack, J. A., & Hutchinson, J. R. (2012). Three-dimensional limb joint mobility in the early tetrapod Ichthyostega. Nature, 486(7404), 523–526.

Polly, P. D. (2007). Limbs in mammalian evolution. Fins into Limbs: Evolution, Development and Transformation, 245–268.

Reilly, S. M., & Elias, J. A. (1998). Locomotion in alligator mississippiensis: kinematic effects of speed and posture and their relevance to the sprawling-to-erect paradigm. The Journal of Experimental Biology, 201 (Pt 18), 2559–2574.

Romer, A. S. (1922). The locomotor apparatus of certain primitive and mammal-like reptiles. American Museum of Natural History.

Romer, A. S. (1924). Pectoral limb musculature and shoulder girdle structure in fish and tetrapods. The Anatomical Record, 27(2), 119–143.

Romer, A. S. (1944). The development of tetrapod limb musculature—the shoulder region of Lacerta. Journal of Morphology.

Romer, A. S., & Price, L. W. (1940). Review of the Pelycosauria. Geological Society of America Special Papers, 28, 1–534.

Romer, A. S. (1956). Osteology of the Reptiles. 772 pp. University of Chicago Press, Chicago.

Romer, A. S. (1967). The Chanares (Argentina) Triassic reptile fauna. III. Two New Gomphodonts, Massetognathus Pascuali and M. Teruggii (Vol. 264). Museum of Comparative Zoology.

Ruta, M., Botha-Brink, J., Mitchell, S. A., & Benton, M. J. (2013). The radiation of cynodonts and the ground plan of mammalian morphological diversity. Proceedings. Biological Sciences / The Royal Society, 280(1769), 20131865.

Simon, W. H. (1970). Scale effects in animal joints. I. Articular cartilage thickness and compressive stress. Arthritis and Rheumatism, 13(3), 244–256.

Stein, B. R. (1981). Comparative Limb Myology of Two Opossums, Didelphis and Chironectes. Journal of Morphology, (169), 113–140.

Stein, B. R. (1986). Comparative limb myology of four arvicolid rodent genera (mammalia, rodentia). Journal of Morphology, 187(3), 321–342.

Sun, A., & Li, Y. (1985). The postcranial skeleton of the late tritylodont Bienotheroides. Vertebrata PalAsiatica, 23(2).

Taylor, B. K. (1978). The anatomy of the forelimb in the anteater (Tamandua) and its functional implications. Journal of Morphology, 157(3), 347–367.

Vaughan, T. A., Ryan, J. M., & Czaplewski, N. J. (2013). Mammalogy. Jones & Bartlett Learning, LLC.

Vickaryous, M. K., & Hall, B. K. (2006). Homology of the reptilian coracoid and a reappraisal of the evolution and development of the amniote pectoral apparatus. Journal of Anatomy, 208(3), 263–285.

Walter, L. R. (1988). Appendicular Musculature in the Echidna, Tachyglossus-Aculeatus (Monotremata, Tachyglossidae). Australian Journal of Zoology, 36(1), 65–81.

Walthall, J. C., & Ashley-Ross, M. A. (2006). Postcranial myology of the California newt, Taricha torosa. The Anatomical Record. Part A, Discoveries in Molecular, Cellular, and Evolutionary Biology, 288(1), 46–57.

Warburton, N. M., Grégoire, L., Jacques, S., & Flandrin, C. (2014). Adaptations for digging in the forelimb muscle anatomy of the southern brown bandicoot (Isoodon obesulus) and bilby (Macrotis lagotis). Australian Journal of Zoology, 61(5), 402–419.

Watson, D. M. (1917). The Evolution of the Tetrapod Shoulder Girdle and Fore-limb. Journal of Anatomy, 52(Pt 1), 1–63.

Witmer, L. M. (1995). The Extant Phylogenetic Bracket and the Importance of Reconstructing Soft Tissues in Fossils. In J. Thomason (Ed.), Functional Morphology in Vertebrate Paleontology.

Zaaf, A., Herrel, A., Aerts, P., & De Vree, F. (1999). Morphology and morphometrics of the appendicular musculature in geckoes with different locomotor habits (Lepidosauria). Zoomorphology, 119(1), 9–22.

